# Multiple classes of spindle-like oscillations reveal stereotyped lifespan dynamics amid individual variability

**DOI:** 10.64898/2026.11.20.768147

**Authors:** Habiba Noamany, Thomas Possidente, Sophie Saremsky, Shuqiang Chen, Ritchie E. Brown, Shaun M. Purcell, Michael Prerau

## Abstract

Sleep spindles, 9–16 Hz transient oscillatory bursts in the sleep electroencephalogram (EEG), are linked to memory consolidation and are altered in aging and disease states. Recent work suggests that traditionally detected spindles are only a subset of a larger class of spindle-like events, and that beyond spindles exist several classes of spindle-like transient oscillatory activity. Together these classes have been shown to comprise individualized signatures of brain activity linked to disease states including schizophrenia and REM behavior disorder. While traditional spindle activity has been studied across the lifespan, the evolution of the full set of spindle-like oscillation classes as a function of age is unknown. To address this major knowledge gap, we characterize the cross-sectional dynamics of multiple classes of spindle-like oscillations in whole-night polysomnography from the Cleveland Family Study (*n* = 704, age: 7–89 y). Unsupervised clustering revealed five classes of spindle-like events in the delta (2–4 Hz), theta (4–6 Hz), theta-alpha (6–9 Hz), sigma-low (9–12 Hz) and sigma-high (12–16 Hz) ranges, each with separate, stereotyped cross-sectional dynamics across the lifespan. Large heterogeneity was observed even within narrow classically defined demographics. Early life (∼10–25 years) is marked by parallel age-related increases in the frequency of the sigma-high and theta-alpha classes and a rapid shift of fast-spindle coupling toward the slow-oscillation peak — timing thought to be critical for memory consolidation. Midlife (∼25–50 years) marks the maximal expression of the delta and sigma-low classes together with an increasingly central-dominant distribution of fast spindles. Late life is defined by progressive uncoupling of lower frequency events from the slow oscillation as well as a leftward shift of higher frequency events away from the peak of the slow oscillation. We further uncover a developmental axis in which slow-oscillation phase coupling decreases in frequency dependence with age. Moreover, we show that the combination of expressed event classes can define phenotypes whose prevalence shifts systematically with age. Together, these provide a robust multi-modal framework for characterizing lifespan dynamics, with wide applications spanning both mechanistic and clinical studies.

## Introduction

Characterizing biological change across the human lifespan is fundamental for understanding the processes underlying development, maturation, and aging, and is critical for tracking health and distinguishing typical from pathological change. Just as pediatric growth charts provide expected trajectories of height and weight^1^, tracking multiple features of brain activity across the lifespan could play an essential role in the early detection of neurodevelopmental and neurodegenerative disorders and for establishing normative reference trajectories of brain function. Lifespan dynamics have been described across multiple domains, including system-wide molecular omics profiles^2^, brain connectivity^3^, and structural “brain charts”^4^. Sleep provides a particularly powerful window into these processes as it is a universal and naturally recurring state involving coordinated autonomic, behavioral, and neural activity across distributed brain circuits^5–9^.

The sleep electroencephalogram (EEG) is a deeply informative and non-invasive approach for characterizing neural activity across the lifespan. Nearly every neurodevelopmental, neurodegenerative, and psychiatric disorder is associated with sleep dysfunction^10–15^, and both sleep architecture and sleep EEG features change systematically with age and disease^5,16–20^. Among these features, sleep spindles, transient oscillatory bursts of 9–16 Hz activity generated by thalamocortical circuitry^10,21,22^, have emerged as a prominent biomarker of brain health. Spindles are the chief marker of stage 2 non-rapid eye movement (NREM) sleep, contribute to sleep-dependent memory consolidation^10,21,23^ and are thought to reflect the functional integrity of thalamocortical circuits^24,25^. Alterations in spindle density, frequency, and timing have been reported in autism^25,26^, schizophrenia^15,27,28^, and Alzheimer’s disease^12,29–32^, and multiple studies have shown clear changes across typical aging^5–8^. While offering many important insights, recent studies have suggested that traditional spindle analyses do not provide a complete picture of the transient oscillatory activity during sleep, thus potentially limiting their power in clinical and mechanistic inference.

Specifically, comparisons of time and time-frequency domain EEG showed that, due to the amplitude bias implicit in standard thresholding, traditionally detected spindles represent only a small morphologically indistinct subset of a larger class of spindle events^33^. Use of the larger class provided both greater statistical power and significantly reduced night-to-night variability in within-subject measurements. Furthermore, emerging evidence suggests that spindles are only one of multiple classes of transient oscillatory activity that occurs across sleep states. Previous case reports and studies have noted alpha bursts around rapid eye movement (REM) sleep^34,35^, oscillations in the delta range in anesthetized rodents^36,37^, and theta bursts in NREM sleep^38,39^. Notably, recent analyses of transient oscillation time–frequency events in humans suggest that sleep spindles represent only a subset of a broader family of “spindle-like” transient oscillatory events spanning the delta, theta, alpha, and sigma frequency ranges^33,40^. The activity from the tens of thousands of spindle-like events across all classes occurring in a single night can be used to construct trait-like neural signatures, shown to be highly individualized yet consistent night-to-night^40^. These signatures have revealed novel robust differences in the transient activity in clinical cohorts including patients with schizophrenia^40^, REM behavior disorder^41^, and fibromyalgia^41^. Together, these observations suggest that neural activity during sleep may be further organized by a coordinated multi-frequency system of transient oscillations.

Thus, while traditional spindle activity over age has been widely studied, the lifespan dynamics of the full extent of spindle and spindle-like oscillations remain unknown. To address this major knowledge gap, we undertake a large-scale population analysis of multi-class transient oscillatory activity during sleep across the human lifespan by analyzing polysomnographic recordings from 704 participants in the Cleveland Family Study^42,43^ spanning 7–89 years. We characterize the population-level organization of spindle-like transient oscillatory events across a broad frequency range during NREM sleep. Using unsupervised clustering, we identify and quantitatively characterize robust classes of spindle-like oscillations in delta (2–4 Hz), theta (4–6 Hz), theta-alpha (6–9 Hz), sigma-low (9–12 Hz), and sigma-high (12–16 Hz) ranges, which form a multi-oscillatory framework for tracking neural dynamics during NREM sleep throughout the lifespan.

## Results

### Data and approach overview

Recent advances in sleep EEG signal processing have provided a quantitative framework for capturing the activity of tens of thousands of spindle-like transient oscillation events across a broad frequency range. The activity from all of these events is then used to construct neural signatures, demonstrated to be highly individualized yet consistent night-to-night^40^. These signatures map the rate of transient oscillatory activity across a frequency range to the power and phase of the slow oscillation (SO), a fundamental sleep rhythm between 0.3 and 1.5 Hz reflecting cortical up- and down-states^44–46^. Slow oscillation power (SO power) can serve as a continuous and objective signal by which spindle activity can be linked to sleep depth^46–48^, providing a high-resolution correlate of neural state during sleep compared to discrete stage classifications^49^. Slow oscillation phase (SO phase) captures the timing of spindles relative to the slow oscillation waveform, and is important given the established relationship between SO/spindle coupling, memory consolidation, and aging^16,50–52^. Consequently, these signatures are called the slow oscillation power histogram (SOPH) and slow oscillation phase histogram (SOΦH), the construction of which is described in detail in Figure 1 and the Methods.

**Figure 1:**
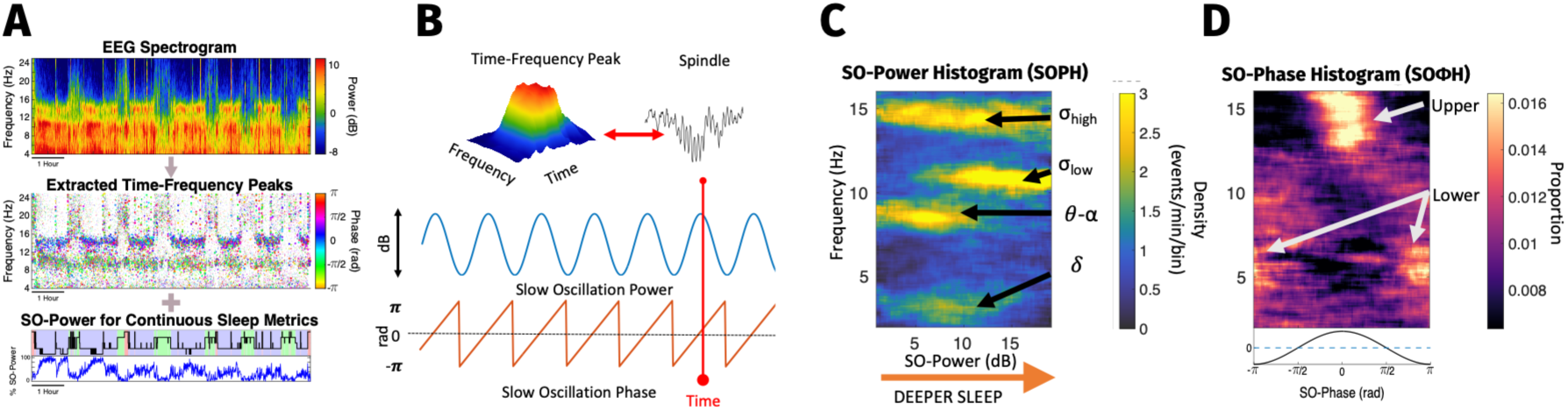
Construction and interpretation of neural signatures of spindle-like activity across sleep. **A.** Thousands of transient time–frequency peaks corresponding to spindle-like oscillatory events are detected across the full night of sleep from the EEG spectrogram (adapted from Stokes et al. 2023^40^). **B.** Each time–frequency peak corresponds to a spindle-like event in the time domain. For each event, features are extracted including central frequency and the slow oscillation (SO) power and phase at the event time. **C.** The Slow Oscillation Power Histogram (SOPH) summarizes events by plotting event frequency (y-axis) as a function of SO power (x-axis; a continuous measure of sleep depth defined as the cumulative power between 0.3 and 1.5 Hz). Color reflects event density for each frequency/SO power bin (number of events/time spent in the SO power defined by that bin, brighter colors = higher density). The SOPH shows how activity changes as a function of frequency and sleep depth, extending beyond conventionally defined sigma-band spindles. High-density regions (modes) indicate individualized classes of spindle-like activity. **D.** The Slow Oscillation Phase Histogram (SOΦH) summarizes events by plotting event frequency (y-axis) as a function of SO phase (x-axis), where each bin is the density of events, normalized by the total number of events at that frequency to reflect the phase-coupling profile for that frequency band. Color reflects the proportion of events at a given frequency that occur at each phase (brighter colors = stronger coupling). The SOΦH reveals frequency-specific phase coupling and distinct modes with similar frequency and phase-coupling characteristics.

This framework builds on the concept that transient oscillatory events in the time domain will appear as peaks (referred to as TF-peaks throughout) in the time–frequency domain, and the finding that canonical spindles represent only a subset of such events^33^. Using an enhanced version of the spectrogram-based approach initially described in Stokes et al.^40^, time–frequency peaks are detected across the full night and a broad 2–16 Hz frequency range (Figure 1A) and event features such as frequency and central time are extracted. From the event’s central time, the associated slow oscillation power and phase can be obtained by mapping that time point onto the slow oscillation power and phase at that time (Figure 1B). The SOPH and SOΦH represent the distribution of event density as a function of frequency and slow oscillation power or phase, respectively. The SOPH characterizes the dynamical structure of spindle-like activity during sleep, showing the density of events (colormap) as a function of event frequency (y-axis) and sleep depth as measured in SO power (x-axis) (Figure 1C). The SOΦH characterizes SO phase coupling structure of transient events by showing the normalized distribution of density (colormap) across each frequency bin (y-axis) as a function of slow-oscillation phase (x-axis) (Figure 1D).

To characterize sleep-mediated transient oscillation dynamics across the lifespan in a cross-sectional population, we analyze the polysomnographic records from 704 participants from the Cleveland Family Study^42,43^ (age range: 7–89 years, 55% female), recorded with a Compumedics E-Series System, using re-referenced C3 and Fpz electrodes (linked mastoid). For each subject, we use the DYNAM-O toolbox^40^ to extract up to tens of thousands of 2–16 Hz spindle-like events during NREM sleep, and to generate slow oscillation power and phase histograms (SOPH and SOΦH) — highly individualized sleep EEG signatures summarizing spindle-like activity with respect to frequency, sleep depth, and phase coupling. Morphologically and phenomenologically similar transient events form high-density regions, or modes, in the parameter space, representing distinct event classes. We quantify the properties (e.g., density, frequency, sleep depth, and phase coupling) of all modes for each individual using a parametric modeling approach in the DYNAM-O toolbox and use unsupervised clustering to group the individually detected modes into different event classes. We then characterize the lifespan dynamics of each event class property as a function of age and electrode and control for several covariates using a regression framework. See Methods for detailed descriptions of the data and quantitative approach.

### Transient event lifespan dynamics reflect trends of neurodevelopment and aging

In order to better characterize neurophysiological changes in transient oscillation activity during development and aging, we show the population mean SO power and phase histograms within 5-year non-overlapping age bins centered at 10 to 75 years at both the central and frontal electrodes (Figure 2).

**Figure 2:**
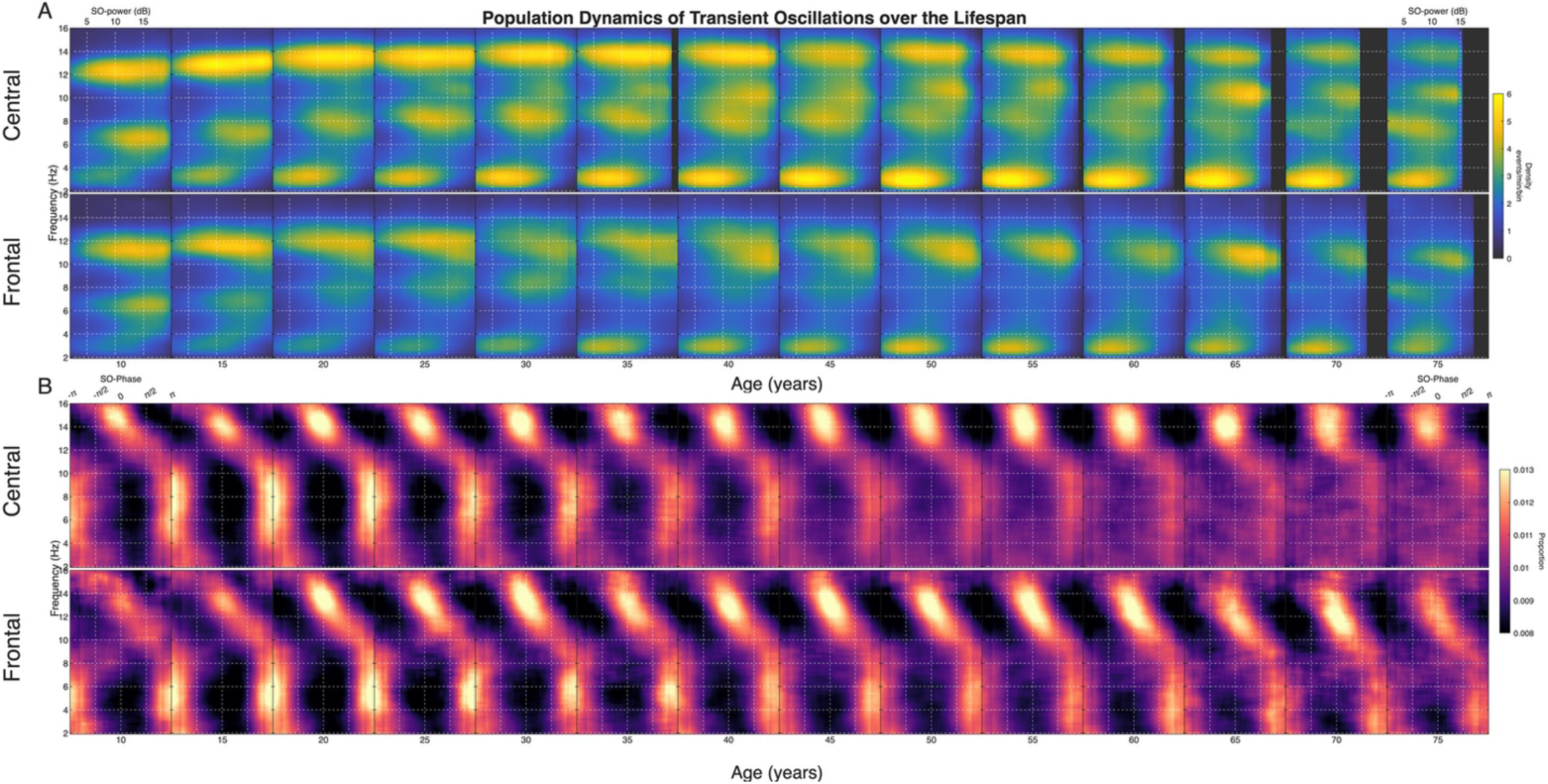
The individualized classes of spindle-like activity form population-level classes that display stereotyped age-related dynamics across the population. To characterize the population-level patterns in the spindle-like classes over age, we computed the SOPHs and SOΦHs for each subject in the Cleveland Family Study (CFS) and averaged them in non-overlapping 5-year age bins, with 682 subjects represented across the displayed 10- to 75-year range (the same set at both frontal and central electrodes). A. The age-binned average SOPHs (top: central electrode, bottom: frontal electrode) from 10 to 75 years show distinct modes of activity that correspond to the different power-mode classes in the delta (2–4 Hz), theta-alpha (6–9 Hz), sigma-low (9–12 Hz), and sigma-high (12–16 Hz) ranges These classes display age-related changes in their center frequency, sleep depth (SO power), and prevalence. Notably, there is an age-related frequency increase in the modes in sigma-high and theta-alpha, the emergence of a sigma-low mode at approximately 30 years, an age-related shift toward earlier sleep depth in theta-alpha, and an age-related increase in delta activity. B. The age-binned average SOΦHs (top: central electrode, bottom: frontal electrode) from 10 to 75 years show two classes with strong phase coupling, an upper class in the fast-sigma range and a broad lower class. These classes also show age-related changes in preferred phase, frequency dependence, and coupling strength. Notably, the upper class shows an age-related decrease in the frequency dependence of phase coupling, together with a shift toward the rising phase, whereas the lower class shows a marked age-related decrease in coupling strength.

The population mean SOPH as a function of age (Figure 2A) reveals distinct lifespan dynamics for different transient oscillation classes in the delta (2–4 Hz), theta-alpha (6–9 Hz), sigma-low (9–12 Hz) and sigma-high (12–16 Hz) ranges, consistent with previous analyses in a small control dataset^40^. Centrally, the sigma-high mode increases in frequency between 10 and 40 years, before plateauing and reducing in density between 40 and 75 years. This is consistent with previous findings of age-related increases in fast spindle frequency^5–8^. Frontally, the sigma-high mode also increases in frequency with age but is lower in frequency compared to the central electrode and is less distinguishable from the sigma-low mode. Centrally, the sigma-low mode is not initially apparent and does not present strongly until about 25–30 years, becoming more prominent at ∼40 years. Frontally, the separation between sigma-high and sigma-low is less distinct, but the sigma-low mode seems to appear earlier frontally, at 15–20 years. The theta-alpha mode shows a striking neurodevelopmental change both centrally and frontally, with a jump in frequency from ∼7–8 Hz between 10–25 years. The theta-alpha mode also becomes more diffuse with aging, which is more pronounced at the frontal electrode. The theta-alpha mode also appears to occur earlier in sleep depth with age, decreasing in SO power from ∼15 dB at 10 years to ∼10 dB by age 50. The delta class of events shows an increase in density with age and has a higher density centrally. Overall, these results suggest that each of these event classes undergoes age-related changes throughout the lifespan, characterized by distinct dynamics as a function of age.

### Aging reshapes SO coupling via upper mode precession, loss of frequency dependence, and lower mode diffusion

The population mean SOΦHs (Figure 2B) also show age-based dynamics. Centrally, the upper mode population mean consistently spans about 12–16 Hz throughout the lifespan. The preferred coupling phase shows a continuous precession, shifting earlier in the SO cycle with age, with the mode centroid moving leftward (toward −π) from just after the SO peak to the rising edge of the SO (∼−π/4 rad). These findings confirm and connect previous observations showing an earlier phase of fast spindle SO coupling in old versus young cohorts^16,54^. Perhaps the most visually apparent change in the central upper mode coupling structure with age is the tilt of the mode itself, which defines the relationship between the transient oscillation frequency and the preferred coupling phase. At the 10-year bin, the mode is highly skewed, with the phase preference shifting earlier in the cycle from the SO falling edge at π/4 to the SO rising edge at −π/4 as oscillation frequency increases from 12 to 16 Hz. This structure suggests a strong dependence between SO coupling and transient oscillation frequency, such that the higher frequency bursts will tend to occur earlier in the SO phase than those with lower frequency. With increasing age, the skew progressively diminishes, such that the mode approaches verticality by the 70-year bin, indicating that all events within the 12–16 Hz range share a similar phase coupling preference. Together, these findings imply that age brings about not only an earlier shift in central coupling preference, but also a progressive loss of frequency dependence, which could serve as a new potential indicator of the aging process. Although a similar leftward shift in the phase coupling of the upper mode is observed frontally, the upper mode remains consistently skewed and thus frequency dependent throughout the lifespan. This suggests that frequency-dependent phase coupling mechanisms may be region specific.

The population mean of the lower phase coupling mode is much broader, spanning ∼4–10 Hz, and showing a coupling preference at the SO trough (±π rad). With increasing age, the lower mode appears to decrease in coupling strength and become more diffuse, such that phase coupling appears near uniform in the 75-year age bin at the central electrode. Frontally, the coupling strength of the lower phase coupling mode appears to decrease at a lower rate compared to the central electrode such that coupling of the lower phase coupling mode is still visible in the 75-year bin.

### Unsupervised clustering identifies five distinct classes of transient oscillatory activity

To move beyond population averages and capture individual heterogeneity, we used the DYNAM-O toolbox to parameterize each subject’s SOPH and extract individual mode properties, including density, frequency, and slow oscillation power and phase. In order to use a more principled quantitative approach to class definition, we perform unsupervised clustering of the parameterized frequency and slow oscillation power of all detected modes, allowing the model to select the optimal number of classes (see Methods). The results of the clustering are shown in Figure 3, displayed as a function of both frequency and sleep depth (Figure 3A,C), and frequency and age (Figure 3B,D). The unsupervised clustering automatically identifies five clusters at both the central and frontal electrodes, corresponding to four previously observed classes: sigma-high, sigma-low, theta-alpha, and delta, as well as an additional class in the theta range. Mean frequency and sleep depth (central/frontal) are 13.6/12.6 Hz and 13.2/12.2 dB for sigma-high, 10.7/10.8 Hz and 15.8/17.8 dB for sigma-low, 7.8 Hz and 11.9/13.1 dB for theta-alpha, 4.9/5.1 Hz and 11.1/11.7 dB for theta, and 3.2/3.1 Hz and 7.4/7.8 dB for delta. The results show that all identified classes, including theta, are well separated in both frequency/SO power and frequency/age projections. The clustering results are used to define the class-specific data in all subsequent mode analyses.

**Figure 3:**
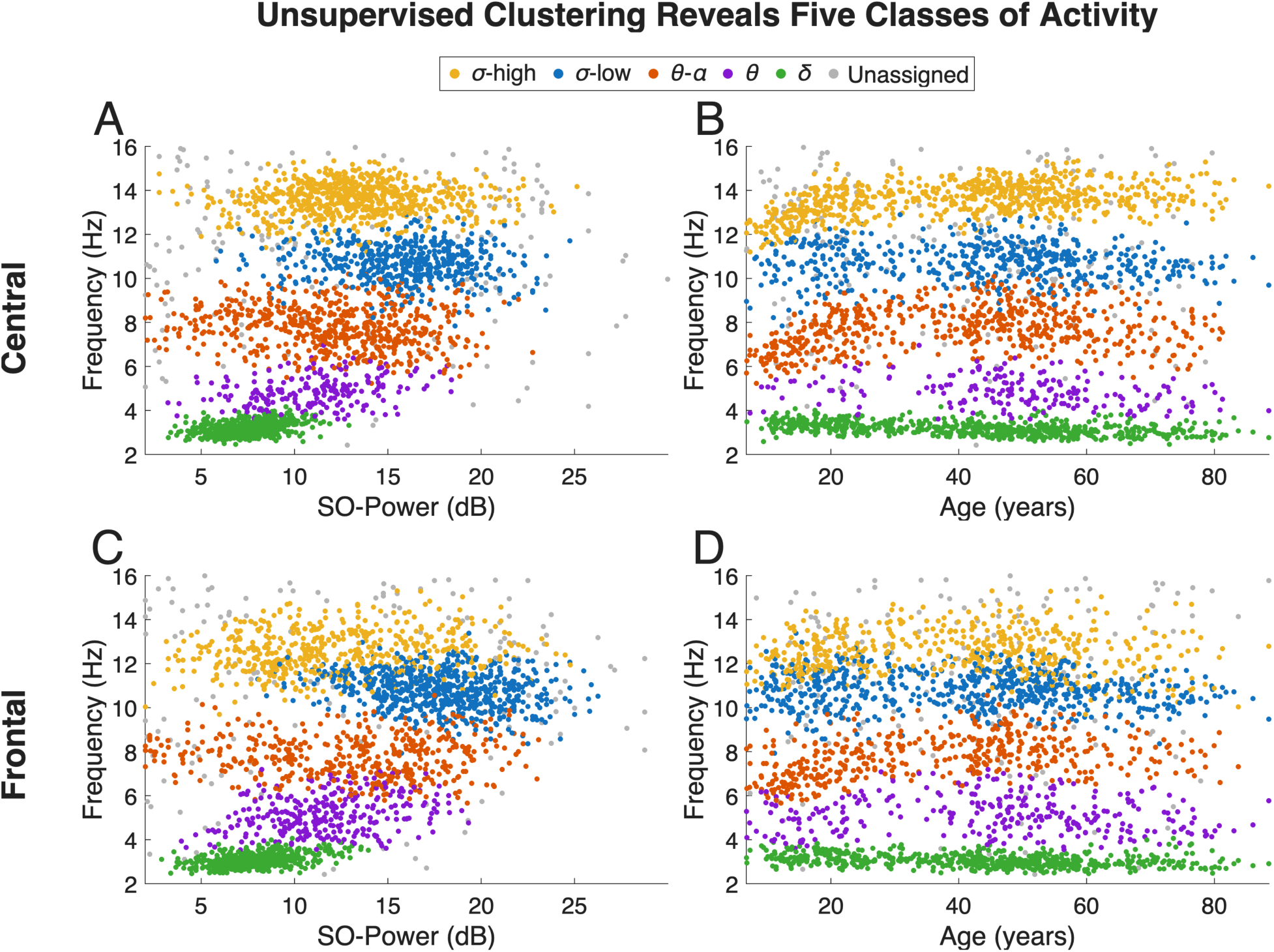
Unsupervised clustering reveals five classes of transient oscillatory activity. Given the presence of multiple classes of activity visually apparent as modes in the SO power histograms (SOPHs), we characterized the distribution of individually detected modes. We used the DYNAM-O toolbox to parameterize the individual SOPHs and identify all the salient modes for each individual histogram and extract the mode properties (center frequency, sleep depth, density, etc.). We then identified how these modes naturally formed classes by clustering all the individually detected modes. We fit a modified Gaussian mixture model to the mode frequency and sleep depth and identified the best number of clusters to fit the data using the Kneedle algorithm (see Methods). Clustering identified five classes of activity (sigma-high, sigma-low, theta-alpha, theta, delta). Modes in gray were assigned to the background. **A,C.** plot the individually detected modes as a function of their center frequency (y-axis) and sleep depth (x-axis) for the central and frontal electrodes and colored by their assigned class (yellow: sigma-high, blue: sigma-low, orange: theta-alpha, purple: theta, green: delta, gray: unassigned)**. B,D.** plot the modes as a function of frequency (y-axis) and age (x-axis). While the modes were clustered based on their frequency and sleep depth, their separation persists when plotted over age.

### Identified event classes are comprised of distinct spindle-like transient events

To characterize the spindle-like events corresponding to the five identified classes, we compute spectrograms and SOPHs for each subject. Figure 4 provides illustrative examples of typical EEG activity described by each event class, derived from the central electrode of four different subjects. For each subject, we show the SOPH and highlight (red rectangle) a mode corresponding to a particular class of event (Figure 4A), individual events in the spectrogram (Figure 4B), along with the bandpass-filtered and clinical time-domain traces (bottom).

**Figure 4:**
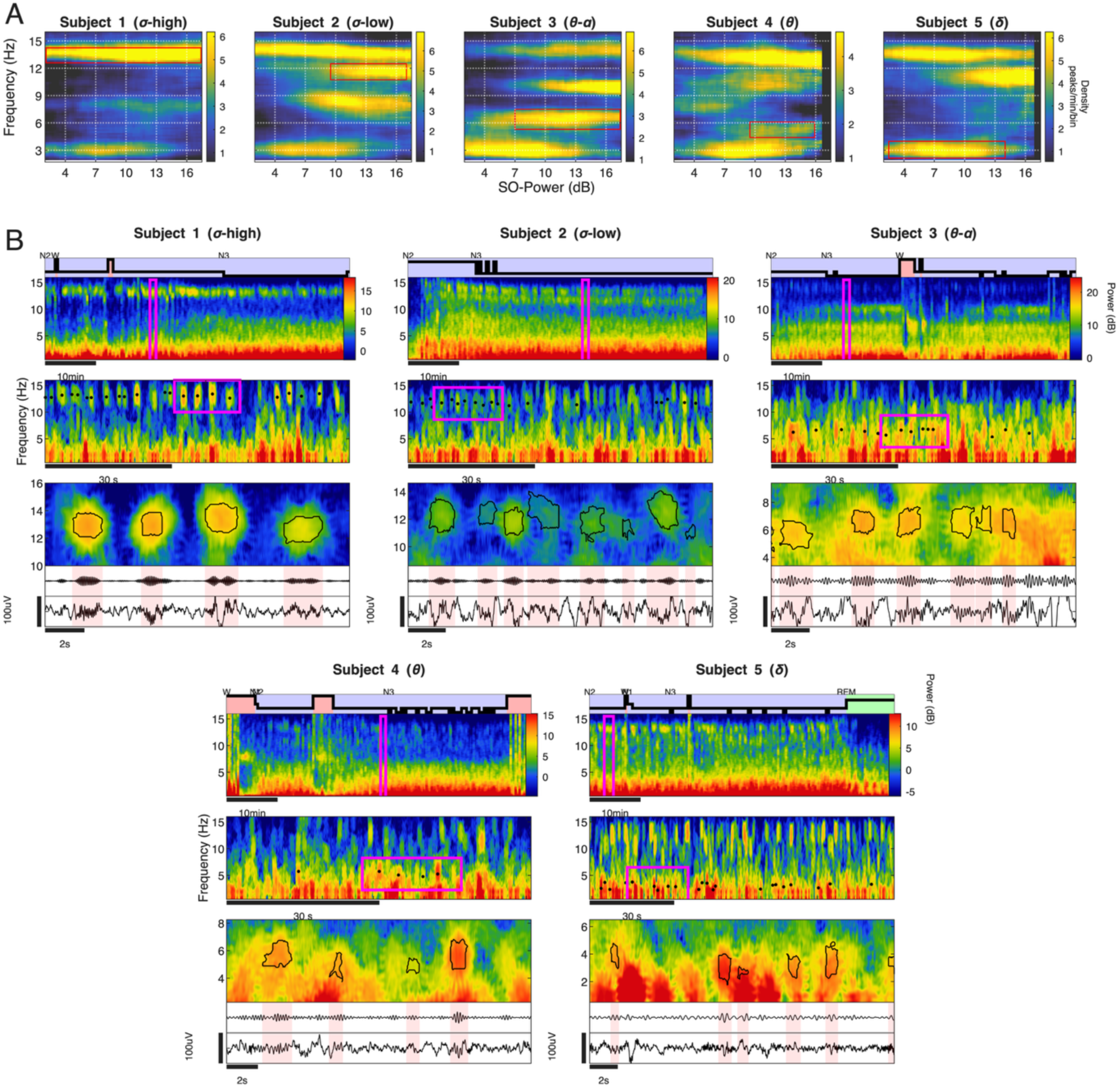
Individuals exhibit spindle-like activity classes beyond the sigma range (G–16 Hz). These individualized classes appear as high-density clusters, or modes of activity on the Slow Oscillation Power Histogram (SOPH) and display distinct properties in the time-frequency and time domains. **A.** Using five different subjects, we highlight each of the five spindle-like classes we observed: the sigma-high class in subject 1, sigma-low in subject 2, theta-alpha in subject 3, theta in subject 4, and delta in subject 5. For each subject, we visualized all the detected spindle-like events from the central electrode using the Slow Oscillation Power Histogram (SOPH), which plots a 2D histogram of the density of all the spindle-like events as a function of their frequency and SO power (a metric of depth of sleep). Spindle-like events with similar frequency and sleep depth form high-density clusters, or modes of activity that represent individualized classes. For each subject, we highlight an individualized spindle-like class as it appears as a distinct mode, localized in frequency and SO power, on the SOPH, indicated with a red rectangle. **B**. For each subject, we show that the events that make up the SOPH modes, or individualized classes, correspond to distinct events in the time-frequency and time domains. For each class, we show example events from that class in three spectrograms with increasing time resolution. In the first spectrogram, we highlight the band of activity corresponding to events in that class. Above the first spectrogram, we display the hypnogram of the scored sleep stages (descending from Wake (W) at the top and NREM3 (N3) at the bottom) in the corresponding time period. In the second spectrogram, we mark the events using black points. In the third spectrogram, with the greatest time resolution, we draw the boundary around the detected events and below it we highlight in pink the corresponding bandpass filtered time trace (top) as well as the unfiltered trace (bottom). The third spectrogram is the average of the 2 spectrograms used in the 2-step TF-peak detection procedure used by DYNAM-O (see Methods). The range of the bandpass filter was determined using the frequency width of each mode.

Subject 1 shows a clear example of sigma-high range activity, encompassing the subset of events typically defined as “fast spindles”^33^. In this subject, these events form a mode in the power histogram that spans approximately 13–14 Hz in frequency, and 2–16 dB in SO power (corresponding to N2 and N3 sleep). As expected, we observe a clear sigma power band at ∼14 Hz in the 1 h spectrogram, which resolves to individual time-frequency peak events (TF-peaks) as the time scale decreases. These TF-peaks correspond directly with ∼1.5 s bursts of oscillatory activity in the bandpass-filtered signal and clinical time-domain traces, consistent with the numerous long-reported descriptions of fast spindles^49,53^.

Subject 2 exhibits clear sigma-low range activity in addition to sigma-high activity, with sigma-low comprising transient events occurring in deeper NREM sleep at a frequency just below sigma-high. The sigma-low mode likely encompasses many of the events defined as “slow spindles” (see Discussion). In this subject, the sigma-low mode occupies the region in the SOPH spanning ∼11–12.5 Hz and 9.5–17 dB in SO power, corresponding with deep N2 to N3 sleep. We observe two bands of activity in the spectrogram in the sigma (9–16 Hz) range, corresponding to sigma-high and sigma-low. These bands resolve into trains of salient TF-peaks at smaller time scales. In this example, we observe interspersed sigma-low and sigma-high events. Overall, the morphology of these slower events in the time and time-frequency domains is highly similar to those in the sigma-high class, differing chiefly in frequency and sleep depth.

Subject 3 illustrates the theta-alpha range class of spindle-like activity, which for this subject appears on the SOPH as a third mode that spans ∼5.5–8 Hz and 7–17.5 dB in SO power. In this case, we observe a power band spanning theta (4–8 Hz) and low alpha (8–9 Hz) activity, which resolves to a set of TF-peaks with a frequency variability higher than the sigma modes. Theta bursts previously reported in NREM^38^ are likely a subset of these events. At the smaller time scales, the increased background/aperiodic EEG activity in this frequency range makes it more difficult to observe the bursts of activity in both the time and time-frequency domains. The relative difficulty of visually identifying theta-alpha class events in the time domain is likely a contributing factor to their under-representation in the literature and to the focus on canonical theta-band power.

Subject 4 provides an example of the theta range class of activity, which appears in this case on the SOPH as a fourth mode that spans ∼4.5-6 Hz and 9.5-16 dB. Compared to the other classes, the theta mode is more diffuse, which likely contributes to its lower prominence in the population average. Although it is diffuse, the clustering results (Figure 3), suggest that there is a separate class of activity in theta that occupies a consistent space in deeper sleep depth, similar to the theta-alpha mode, but at a lower frequency. Correspondingly, a diffuse power band can be seen in the spectrogram at ∼4.5-6Hz, but it is not as distinct as the other classes, likely due to its proximity to the high-power slow waves as well as the higher frequency theta-alpha activity in this subject. This activity resolves into TF peaks, but, as with the theta-alpha events, its low frequency and high background activity make the bursts difficult to observe in the time domain. The challenge is even greater here because these events occur at lower frequencies than the theta-alpha events.

Subject 5 provides an example of the delta range class of activity, which appears in this case as a 2–4 Hz and 4–14 dB mode on the SOPH. Unlike the other classes, the delta events do not form an easily identifiable band of activity on the spectrogram, likely due to the steep slope of the aperiodic EEG at low frequency and the high power of slow waves. At smaller time scales, however, salient TF-peaks are visible in the delta range within the spectrogram. Moreover, these delta-class TF-peaks have a distinct time/time-frequency morphology from the lower-frequency slow-wave activity. Specifically, the delta-class TF-peaks appear as narrow peaks in the spectrogram, whereas slow waves appear as broadband activity emanating from lower frequencies. However, the delta-class events are difficult to see in the time domain given that they may consist of only three or four oscillation cycles.

In addition to characterizing transient oscillations in terms of frequency and sleep depth, the SOΦH characterizes their activity in terms of frequency and slow oscillation phase coupling. As previously noted in the literature^40^, we observe primary phase coupling modes that span the frequency range of multiple event classes (Figure 1). Specifically, there exists an upper mode with a frequency range of approximately 12–16 Hz, exhibiting phase coupling preference near the peak of the slow oscillation (0 rad). Additionally, there is a broader, lower phase coupling mode with a frequency range of approximately 4–10 Hz, characterized by phase preference at the trough of the slow oscillation (∼−π rad).

### Distributions of transient event class properties show significant changes across the lifespan

While the population mean histograms identify age dynamics in the transient event classes, aggregate measures mask individual variability and cannot separate changes of density versus changes of prevalence. Moreover, the SOPHs are highly individualized in nature and exhibit substantial heterogeneity, even within control cohorts^40^, so it is crucial to account for variability in mode features at the individual subject level. We first plot the parameterized features of every detected mode in 5-year age bins (Figure 5A). Each point is one mode from one subject, shown by frequency (y-axis) and sleep depth/SO power (x-axis) for central (top) and frontal (bottom) electrodes. Point size shows mode density, and color shows the preferred SO coupling phase at the mode centroid. Despite clear variability between subjects, the class separation and lifespan trends seen in the population mean (Figure 2) remain clearly visible.

**Figure 5:**
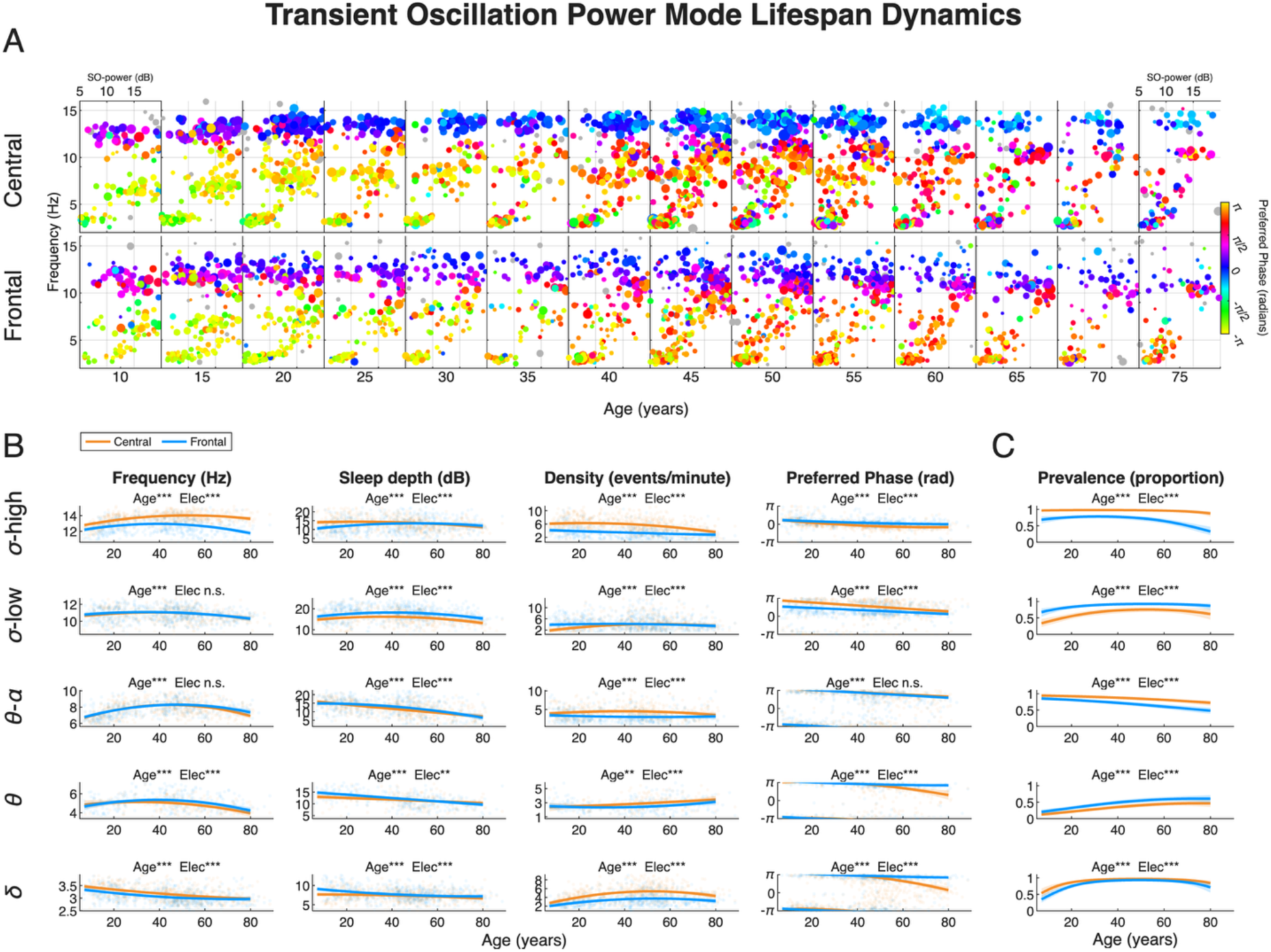
Multiple classes of spindle-like events show significant age-based dynamics over the lifespan. For each participant, frontal and central SOPHs were quantified by parameterizing all modes observed. A. shows the mode-quantified equivalent of Figure 3A, with the detected modes in the central (upper panel) and frontal (lower panel) electrodes plotted as a function of frequency and sleep depth (SO power) in 5-year age bins (size = mode density, color = mode phase preference, excluded modes marked in gray). This visualization shows the natural clustering in multiple dimensions between the different classes, which remain visually separable over age and show similar age-related dynamics observed in the average SOPHs in Figure 3A. B. After categorizing into sigma-high, sigma-low, theta-alpha, theta, and delta modes using clustering, each class feature (frequency, SO power, preferred phase, and density) was fit to age with a subject-level mixed-effects regression matched to that feature — a linear (Gaussian) model for frequency, SO power, and density, and a circular (von Mises) model for preferred phase — adjusting for relevant covariates (sex, electrode, race, AHI, and BMI) and allowing the age dynamics to differ by electrode; the polynomial order in age was selected by AIC (see Methods). The lines represent the model fits for the central (orange) and frontal (blue) electrodes and the gray points represent the individual feature values for all detected modes in that class. Age-related trajectories (indicated by significant polynomial age terms) differed across classes, features, and electrode, indicating that the classes undergo distinct changes in their frequency, sleep depth, density, and preferred phase across the lifespan with spatial differences across electrodes. C. Prevalence of each class across the lifespan, shown per electrode (central = orange, frontal = blue), from a binomial logistic regression of per-subject class expression on age (adjusted for electrode, sex, race, AHI, and BMI), with the fitted curve and a 95% confidence band obtained from a family-level bootstrap stratified by family mean age. Class prevalence showed significant age-related changes and also differed across electrode with some classes increasing in prevalence at different rates (indicated by different age terms) such as central sigma-low, theta, and delta and others decreasing with age such as the frontal sigma-high and theta-alpha. Stars indicate the significance level for an omnibus Wald test of all age-related terms and for all electrode-related terms.

### Ǫuantifying dynamics of individualized transient oscillatory class features throughout the lifespan

For each feature of each mode class, we fit a mixed-effects regression on a polynomial function of age, with order selected by minimum Akaike information criterion (AIC). Fixed effects include electrode, sex, race, apnea–hypopnea index (AHI), and body-mass index (BMI), plus interactions of the age terms with sex and electrode. A per-subject random intercept accounts for correlation between a subject’s frontal and central observations. A family-level random intercept accounts for correlation among relatives. The phase models account also account for correlation among relatives (See Methods), To estimate the prevalence of each class, we model the probability that a subject expresses the class on a given electrode using logistic regression on the same polynomial in age (order by AIC), with sex, electrode, race, AHI, and BMI as covariates. Prevalence confidence intervals and significance tests are estimated via a bootstrap that resamples whole families, accounting for relatives and for each subject’s two electrodes (see Methods). Figure 5B shows the estimated age dynamics (regression lines) for the key mode features (frequency, SO power, density, and preferred phase) at the central (orange) and frontal (blue) electrodes, with each column corresponding to a feature and each row to a class. Figure 5C shows the estimated age dynamics (regression lines) for the prevalence of the five classes. The full model statistics for each SOPH class are summarized in Table 1. In the following sections, we describe the observed age and electrode effects for each class.

**Table 1:** Lifespan statistics for power modes (SO power histogram, SOPH), organized by mode. For each power-mode class, each feature (frequency, SO power, density and preferred phase) was regressed on age, pooling central (C3) and frontal (Fpz) electrodes with electrode as a within-subject factor. All models included a polynomial in age (order 0–2, selected by AIC), sex, electrode, race, AHI and BMI, age × sex and age × electrode interactions, a per-subject random intercept, and a family random intercept (family-clustered standard errors for preferred phase). Age is an omnibus Wald test that tests all polynomial age terms together with their sex and electrode interactions. Sex and Electrode test each main effect together with its age interactions. Race, AHI and BMI test single coefficients. Prevalence (the proportion of subjects expressing the class) was modeled by binomial logistic regression with a family-level bootstrap stratified by family mean age (5,000 resamples; n = 704). For all rows: n, number of subjects; R², variance explained by the fixed effects (marginal R²). R² is not reported for prevalence (–), whose binary outcome has no directly comparable R².

|  | *P* value |  |  |  |  |  | R<sup>2</sup> |
|---|---|---|---|---|---|---|---|
| **Mode / feature** | **Age** | **Sex** | **Electrode** | **Race** | **AHI** | **BMI** |  |
| **Sigma-high** (n = 692) |  |  |  |  |  |  |  |
| Frequency | 1.9 × 10<sup>−33</sup> | 7.7 × 10<sup>−4</sup> | 3.5 × 10<sup>−128</sup> | 0.039 | 0.014 | 0.20 | 0.41 |
| SO power | 5.0 × 10<sup>−8</sup> | 0.26 | 8.6 × 10<sup>−12</sup> | 0.0016 | 0.0033 | 0.23 | 0.07 |
| Density | 1.8 × 10<sup>−21</sup> | 0.26 | 3.2 × 10<sup>−84</sup> | 0.72 | 6.3 × 10<sup>−4</sup> | 0.20 | 0.33 |
| Preferred phase | 3.7 × 10<sup>−23</sup> | 0.51 | 1.1 × 10<sup>−29</sup> | 0.65 | 0.95 | 0.44 | 0.23 |
| Prevalence | 1.6 × 10<sup>−7</sup> | 0.35 | 3.5 × 10<sup>−31</sup> | 0.25 | 0.41 | 0.13 | – |
| **Sigma-low** (n = 663) |  |  |  |  |  |  |  |
| Frequency | 3.9 × 10<sup>−5</sup> | 0.26 | 0.52 | 4.5 × 10<sup>−4</sup> | 0.93 | 0.16 | 0.06 |
| SO power | 6.2 × 10<sup>−11</sup> | 0.027 | 5.1 × 10<sup>−34</sup> | 0.062 | 0.74 | 0.53 | 0.15 |
| Density | 2.8 × 10<sup>−11</sup> | 0.63 | 9.9 × 10<sup>−14</sup> | 7.0 × 10<sup>−8</sup> | 8.1 × 10<sup>−5</sup> | 0.57 | 0.11 |
| Preferred phase | 1.7 × 10<sup>−12</sup> | 0.20 | 2.1 × 10<sup>−27</sup> | 0.0035 | 0.15 | 0.33 | 0.21 |
| Prevalence | 2.1 × 10<sup>−6</sup> | 0.061 | 4.3 × 10<sup>−22</sup> | 0.0016 | 0.28 | 0.39 | – |
| **Theta-alpha** (n = 620) |  |  |  |  |  |  |  |
| Frequency | 6.9 × 10<sup>−34</sup> | 0.28 | 0.065 | 0.0090 | 0.29 | 0.35 | 0.26 |
| SO power | 5.6 × 10<sup>−35</sup> | 0.93 | 4.5 × 10<sup>−6</sup> | 0.056 | 0.022 | 0.93 | 0.21 |
| Density | 2.7 × 10<sup>−6</sup> | 4.0 × 10<sup>−5</sup> | 2.5 × 10<sup>−59</sup> | 0.71 | 0.0050 | 0.0054 | 0.16 |
| Preferred phase | 8.7 × 10<sup>−26</sup> | 0.0064 | 0.28 | 0.96 | 0.64 | 0.78 | 0.20 |
| Prevalence | 3.9 × 10<sup>−11</sup> | 0.0024 | 1.5 × 10<sup>−16</sup> | 0.14 | 0.36 | 0.48 | – |
| **Theta** (n = 368) |  |  |  |  |  |  |  |
| Frequency | 1.1 × 10<sup>−8</sup> | 0.046 | 2.7 × 10<sup>−4</sup> | 0.013 | 0.83 | 0.64 | 0.14 |
| SO power | 5.2 × 10<sup>−16</sup> | 0.61 | 0.0036 | 0.0031 | 0.16 | 0.39 | 0.24 |
| Density | 0.0057 | 0.42 | 3.3 × 10<sup>−6</sup> | 0.15 | 0.045 | 0.32 | 0.09 |
| Preferred phase | 2.6 × 10<sup>−13</sup> | 0.36 | 4.9 × 10<sup>−4</sup> | 0.60 | 0.78 | 0.31 | 0.17 |
| Prevalence | 3.6 × 10<sup>−11</sup> | 0.055 | 3.7 × 10<sup>−8</sup> | 0.043 | 0.26 | 0.12 | – |
| **Delta** (n = 662) |  |  |  |  |  |  |  |
| Frequency | 5.0 × 10<sup>−25</sup> | 0.41 | 1.0 × 10<sup>−19</sup> | 0.61 | 0.86 | 0.92 | 0.18 |
| SO power | 1.3 × 10<sup>−9</sup> | 0.31 | 2.1 × 10<sup>−13</sup> | 0.71 | 0.085 | 0.55 | 0.08 |
| Density | 2.0 × 10<sup>−22</sup> | 0.63 | 8.0 × 10<sup>−136</sup> | 0.51 | 0.25 | 0.20 | 0.27 |
| Preferred phase | 2.3 × 10<sup>−28</sup> | 0.74 | 1.1 × 10<sup>−10</sup> | 0.60 | 0.50 | 0.43 | 0.16 |
| Prevalence | 1.7 × 10<sup>−19</sup> | 0.10 | 2.3 × 10<sup>−7</sup> | 0.36 | 0.56 | 0.065 | – |

### Sigma-high events increase in frequency, shift toward the SO rising phase, and become centrally dominant with age

The lifespan dynamics of the sigma-high mode are shown in the first row of Figure 5B, and the summary model statistics are reported in Table 1. The models for frequency, sleep depth, density, and preferred phase, as well as for the prevalence (proportion of total subjects expressing a mode) of sigma-high, all have highly significant effects of age and electrode (p < 0.001). Centrally, the sigma-high frequency increases from just below 13 Hz at 10 years to a peak near 14 Hz around 60 years before a slight decline, while frontally it increases from about 12 Hz to about 13 Hz and remains below central across the lifespan. The significant electrode effect reflects the consistently lower frontal frequency, which is consistent with previous studies of fast spindle frequency at the central electrode^5–8^. For sleep depth, the central sigma-high events occur progressively earlier in NREM with age, with the SO power decreasing steadily from about 15 dB at 10 years to about 12.5 dB at 80 years, whereas frontally the events occur later in NREM with age, with the SO power increasing from about 10 dB at 10 years, reaching about 14 dB by 60 years. The density is higher centrally and decreases with age. Centrally, the density decreases from about 6 events/min at 10 years to about 4.5 events/min at 80 years. Frontally, the age-related decrease is much more modest, from about 4 events/min at 10 years to about 3.5 events/min at 80 years. This decline in density is consistent with previously reported decreases in fast spindle density with age^5,22^. Centrally, the preferred SO phase of the sigma-high events shifts steadily from the falling phase of the SO at about π/4 rad at 10 years, crossing the SO peak (0 rad) near 35 years, and reaching about −π/7 rad on the rising phase by 80 years, with the precession leveling off after about 60 years. The frontal preferred phase follows a parallel precession but remains more positive than central at every age, running from about π/3 rad at 10 years to about −0.16 rad by 80 years. This leftward shift in phase coupling with age is consistent with previously reported shifts in the phase coupling of fast spindles at the central electrode^16,54^.

The sigma-high mode is nearly universal centrally, expressed by about 99% of individuals throughout life, whereas frontally the proportion remains at about 91% from 10–45 years but begins to decrease at 45 years, reaching about 26% by 80 years, indicating that the sigma-high mode becomes increasingly central-dominant with age (Figure 5C).

### Sigma-low events emerge in early-life (10-25 years) and become increasingly prevalent centrally

The lifespan dynamics of the sigma-low mode are illustrated in the second row of Figure 5B, and the model statistics are summarized in Table 1. The models for frequency, sleep depth, density, and preferred phase, as well as for the proportion of subjects expressing the sigma-low mode, all have a highly significant effect of age (p < 0.001) and all except for frequency also have a significant effect of electrode. The sigma-low frequency increases slightly with age, rising from about 11 Hz at 10 years to about 11.3 Hz by 40 years, after which it begins to decrease back to 11 Hz, with no significant effect of electrode. For sleep depth, the sigma-low events occur at progressively higher SO power, that is, deeper in NREM, with increasing age, rising from about 14.6 dB at 10 years to about 15.0 dB at 80 years centrally and from about 15.6 dB to about 16.4 dB frontally. The significant electrode effect reflects the consistently deeper frontal sleep depth, about 1 dB above central across the lifespan. The density shows different dynamics across electrodes. Centrally, it increases markedly with age, rising from about 2.2 events/min at 10 years to a peak near 4.4 events/min around 60 years, while frontally the density stays relatively stable at about 4 events/min throughout. Centrally, the preferred SO phase of the sigma-low events shifts leftward with age from near the SO trough at about 2.97 rad at 10 years to about 1.5 rad by 80 years. Frontally, the preferred phase follows a parallel leftward shift but remains consistently closer to the SO peak than central, moving from about 2π/3 rad (2.09 rad) at 10 years to about π/5 rad (0.63 rad) by 80 years.

Centrally, the sigma-low mode becomes more prevalent with age, rising from about 30% of individuals at 10 years to about 65% by 40 years (Figure 5C). Frontally, the sigma-low is more prevalent and has more consistent expression across age, expressed by about 70% of 10-year-olds and reaching about 90% by about 25 years.

### Theta-alpha events have an early-life frequency increase and shift to lighter sleep with age

The lifespan dynamics of theta-alpha modes are illustrated in the third row of Figure 5B and the model statistics are summarized in Table 1. The models for frequency, sleep depth, density, and preferred phase, as well as for the proportion of subjects expressing the theta-alpha mode, all have a highly significant effect of age (p < 0.001) and all except for frequency and preferred phase also have a significant effect of electrode. The frequency increases most in early life, from ∼7 Hz at 10 years to ∼7.8 Hz by ∼25 years, then increases more slowly to ∼8 Hz by 40 years, after which the frequency begins to slightly decrease, with no significant effect of electrode. For sleep depth, the theta-alpha events occur progressively earlier in NREM with age, with a large, steady decrease in the SO power from ∼16 dB at 10 years to ∼8 dB at 80 years, with the frontal events starting at a slightly lower sleep depth of ∼15 dB and decreasing at a slightly lower rate. The density is higher centrally compared to frontally and shows different age-related dynamics. Centrally, the density increases from about 4 events/min at 10 years to ∼5 events/min by 40 years, after which it begins to decrease. Frontally, the density slightly decreases from ∼3.5 events/min at 10 years to ∼3 events/min by 80 years. The preferred SO phase of the theta-alpha events shifts leftward with age from near the SO trough at 10 years to about 2 rad by 80 years, with no significant effect of electrode. Theta-alpha events are more common centrally than frontally and become less prevalent with age, declining centrally from approximately 95% to 80% of the population between 10 and 80 years, and frontally from 82% to 55% over the same range.

### Theta events have an early-life frequency increase and become more prevalent and occur in lighter sleep with age

The lifespan dynamics of the theta modes are illustrated in the fourth row of Figure 5B and the model statistics are summarized in Table 1. The models for frequency, sleep depth, density, and preferred phase, as well as for the proportion of subjects expressing the theta mode, all have a highly significant effect of age and electrode (p < 0.001), except for density which has a significant effect of age (p<0.01) and sleep depth, which has a significant effect of electrode (p<0.01). The frequency of the theta events is slightly higher frontally and increases from ∼5 Hz at both electrodes at 10 years to ∼5.4 Hz centrally and ∼5.6 Hz frontally by 30 years, after which it begins to decrease, reaching ∼4.5 Hz frontally and ∼4.2 Hz centrally by 80 years. For the sleep depth, the theta events become earlier in sleep depth, with a more pronounced decrease in SO power at the frontal electrode compared to the central, decreasing from about 15 dB at 10 years to about 11 dB at 80 years frontally and 13.5 dB to 12.5 dB centrally. The density increases with age, at a faster rate centrally, starting from ∼2.5 events/min at both electrodes at 10 years and reaching about 4 events/min by 80 years centrally and 3.5 events/min frontally. The preferred phase shifts leftward with age, from the trough of the SO (−*π*) at 10 years toward the later part of the falling phase at 4π /5 rad by 80 years, with the frontal events having a preferred phase slightly closer to the trough of the SO. The theta events increase in prevalence with age and are more prevalent frontally, with ∼12% of individuals at 10 years centrally (∼25% frontally) to ∼50% by 80 years centrally (65% frontally).

### Delta events decrease in frequency and increase in density and prevalence from childhood to midlife

The lifespan dynamics of the delta modes are illustrated in the fifth row of Figure 5B and the model statistics are summarized in Table 1. The models for frequency, sleep depth, density, and preferred phase, as well as for the proportion of subjects expressing the delta mode, all have a highly significant effect of age and electrode (p < 0.00) . The frequency of the delta events is higher centrally and decreases with age from about 3.5 Hz at 10 years to about 3.1 Hz by 80 years centrally. Frontally, the frequency is slightly lower at about 3.4 Hz at 10 years but decreases more slowly, reaching about 3.1 Hz frontally. For the sleep depth, the delta events become slightly earlier in sleep depth with increased age, with the frontal shift being more pronounced. Frontally, the SO power decreases from about 9.5 dB at 10 years to about 7.5 dB by 80 years. Centrally, the SO power decreases from about 7.5 dB at 10 years to about 6.5 dB at 80 years. The density is higher centrally and increases from about 3 events/min at 10 years to about 5 events/min by 45 years, after which it begins to slightly decrease. Frontally, the density increase is more modest, increasing from about 2 events/min at 10 years to about 2.7 events/min at 45 years, after which it begins to slightly decrease. The preferred phase of the delta events closely mirrors that of the theta class, likewise shifting leftward with age from the SO trough (−π) at 10 years toward the late falling phase (∼4π/5 rad) by 80 years, with frontal events remaining slightly closer to the trough. Delta events increase in prevalence from early to midlife, rising from ∼62% at age 10 to 87% by age 40 at the central electrode. Frontal prevalence is lower at age 10 (∼37%) but approaches central prevalence by age 40.

### Distributions of individualized phase coupling mode features group into classes that change over the lifespan

To quantify the dynamics of phase-coupling modes, we follow a similar approach to that used for the SO power histograms. Using the DYNAM-O toolbox, we detect each subject’s phase-coupling modes and extract their features (frequency, preferred phase, coupling strength, etc.). We then classify each mode according to its peak frequency as either lower (4–10 Hz) or upper (10–16 Hz). Figure 6A characterizes trends and variability in the two modes (upper and lower) observed in the SOΦHs using an approach similar to that used for the event classes in the SOPHs. Each point represents a single phase mode from a single subject for the central (top) and frontal (bottom) electrode, plotted as a function of frequency (y-axis) and SO phase (x-axis). The size of each point corresponds to the phase coupling strength, while the color indicates the angle of the tilt of the mode, representing the frequency dependence of the coupling. Similar to the average SOΦH, we observe distinct clusters emerge from the individually detected phase coupling modes, corresponding to the upper and lower phase-coupling classes observed in the SOΦH population mean (Figure 2B). The dynamics of the distributions of the two clusters of modes also appear to correspond to those observed in the SOΦH population mean.

**Figure 6:**
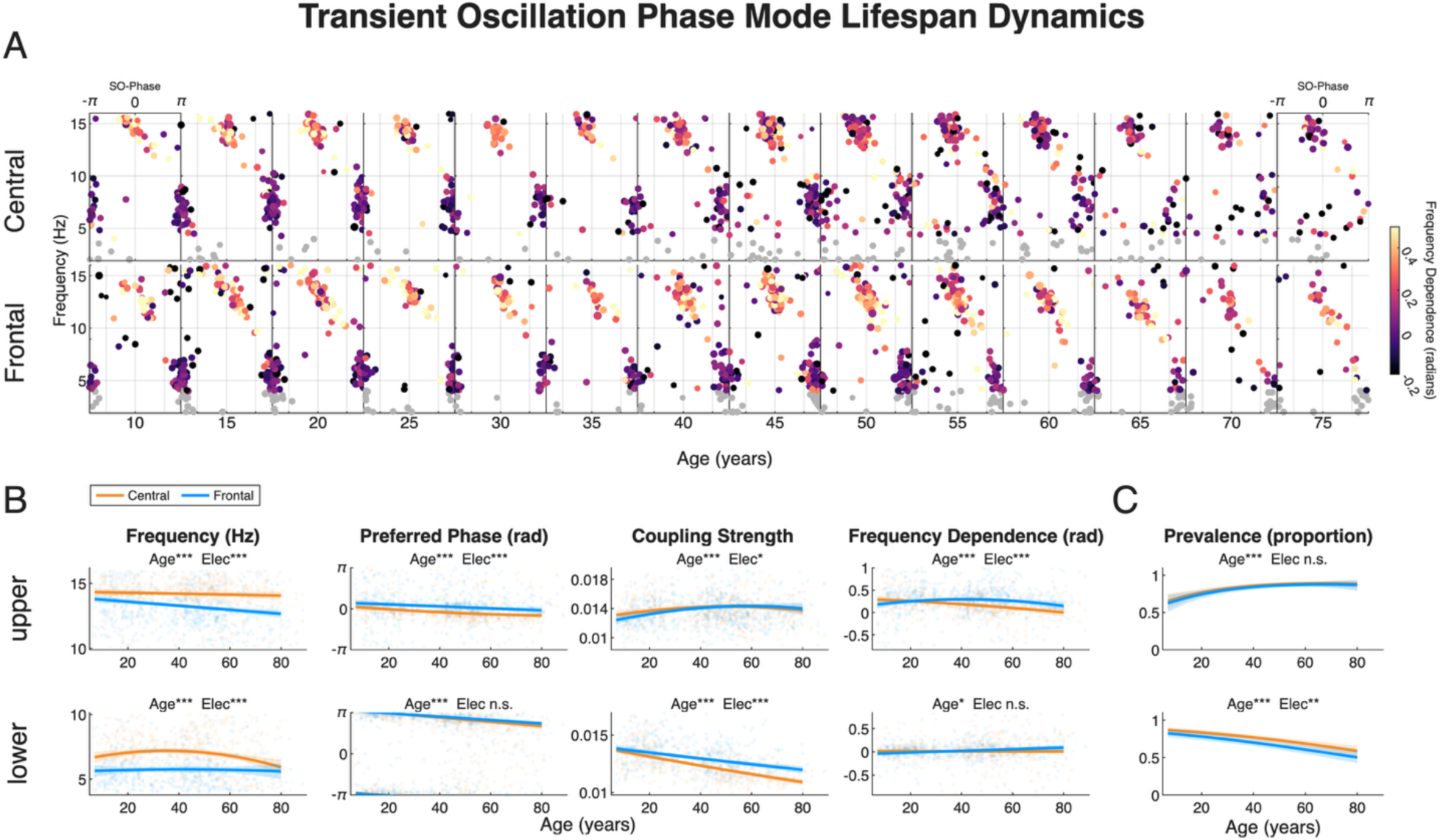
Two classes of phase-coupled transient oscillations show significant age-based dynamics over the lifespan. For each participant, frontal and central SOΦHs were quantified by parameterizing all modes observed. A. shows the mode-quantified equivalent of Figure 3B, with the detected modes in the central (upper panel) and frontal (lower panel) electrodes plotted as a function of frequency and SO phase in 5-year age bins (size = coupling strength, color = mode frequency dependence, excluded modes marked in gray). This visualization shows the natural clustering in multiple dimensions between the different classes, which remain visually separable over age and show similar age-related dynamics observed in the average SOΦHs in Figure 3B. B. After categorizing into upper and lower modes by frequency, each class feature (frequency, preferred phase, coupling strength, and frequency dependence) was fit with a subject-level mixed-effects regression matched to that feature: a linear (Gaussian) model for frequency and frequency dependence, a circular (von Mises) model for preferred phase, and a Gamma (log link) model for coupling strength, adjusting for sex, electrode, race, AHI, and BMI and allowing the age dynamics to differ by electrode. The polynomial order in age was selected by AIC (see Methods). The lines represent the model fits for the central (orange) and frontal (blue) electrodes and the gray points represent the individual feature values for all detected modes in that class. Age-related trajectories (indicated by significant polynomial age terms) differed across classes, features, and electrode, indicating that the classes undergo distinct changes in their frequency, preferred phase, coupling strength, and frequency dependence across the lifespan with spatial differences across electrodes. C. Prevalence of each class across the lifespan, shown per electrode (central = orange, frontal = blue), from a binomial logistic regression of per-subject class expression on age (adjusted for electrode, sex, race, AHI, and BMI), with the fitted curve and a 95% confidence band obtained from a family-level bootstrap stratified by family mean age. Both the upper and lower phase coupling classes showed significant age-related changes in prevalence, with the lower mode showing a marked decrease in prevalence with age. Stars indicate the significance level for an omnibus Wald test of all age-related terms and for all electrode-related terms.

### Ǫuantifying dynamics of individualized phase coupling mode features throughout the lifespan

Similar to the SOPH modes, we quantify the changes in the features of the phase coupling classes with age by fitting a mixed-effects regression of the feature as a polynomial function of age, with main effects of sex, electrode, race, AHI, and BMI, interactions of the age terms with both sex and electrode, a per-subject random intercept, and family structure as above. The full model statistics for each SOΦH class are summarized in Table 2. In the following sections, we describe the observed age and electrode effects for each class.

**Table 2:** Lifespan statistics for phase modes (SO phase histogram, SOΦH), organized by mode. For each phase-mode class (upper, lower), each feature (frequency, preferred phase, coupling strength and frequency dependence) was regressed on age, pooling central (C3) and frontal (Fpz) electrodes with electrode as a within-subject factor. Model structure and covariates were identical to those in Table 1. Age is an omnibus Wald test that tests all polynomial age terms together with their sex and electrode interactions. Sex and Electrode test each main effect together with its age interactions. Race, AHI and BMI test single coefficients. Prevalence (the proportion of subjects expressing the class) was modeled by binomial logistic regression with a family-level bootstrap stratified by family mean age (5,000 resamples; n = 704). For all rows: n, number of subjects; R², variance explained by the fixed effects (marginal R²). R² is not reported for prevalence (–), whose binary outcome has no directly comparable R².

|  | ***P* value** |  |  |  |  |  | **R<sup>2</sup>** |
|---|---|---|---|---|---|---|---|
| **Mode / feature** | **Age** | **Sex** | **Electrode** | **Race** | **AHI** | **BMI** |  |
| **Upper** (n = 663) |  |  |  |  |  |  |  |
| Frequency | 2.0 × 10<sup>−7</sup> | 5.8 × 10<sup>−4</sup> | 1.2 × 10<sup>−40</sup> | 0.38 | 0.062 | 0.35 | 0.15 |
| Preferred phase | 2.2 × 10<sup>−7</sup> | 0.16 | 4.7 × 10<sup>−23</sup> | 0.47 | 0.71 | 0.55 | 0.08 |
| Coupling strength | 2.5 × 10<sup>−5</sup> | 2.1 × 10<sup>−5</sup> | 0.042 | 0.12 | 0.021 | 0.83 | 0.05 |
| Frequency dependence | 9.3 × 10<sup>−8</sup> | 0.10 | 2.3 × 10<sup>−12</sup> | 0.68 | 0.68 | 0.42 | 0.06 |
| Prevalence | 7.7 × 10<sup>−7</sup> | 0.037 | 0.30 | 0.92 | 0.68 | 0.43 | – |
| **Lower** (n = 632) |  |  |  |  |  |  |  |
| Frequency | 5.9 × 10<sup>−4</sup> | 0.89 | 9.2 × 10<sup>−45</sup> | 0.063 | 0.033 | 0.079 | 0.19 |
| Preferred phase | 4.4 × 10<sup>−19</sup> | 0.69 | 0.29 | 0.32 | 0.75 | 0.091 | 0.10 |
| Coupling strength | 3.2 × 10<sup>−40</sup> | 0.084 | 2.3 × 10<sup>−24</sup> | 0.79 | 0.13 | 0.76 | 0.28 |
| Frequency dependence | 0.031 | 0.97 | 0.051 | 0.15 | 0.30 | 0.47 | 0.01 |
| Prevalence | 1.2 × 10<sup>−9</sup> | 0.63 | 0.0014 | 0.055 | >0.99 | 0.90 | – |

### The upper phase-coupling mode shifts toward the SO rising phase, with a loss of central frequency dependence

The lifespan dynamics of upper phase coupling modes are illustrated in the first row of Figure 6B and the model statistics are summarized in Table 2. The models for frequency, preferred phase, coupling strength, and frequency dependence, as well as for the proportion of subjects expressing the upper coupling mode, all have a highly significant effect of age (p < 0.001). All except prevalence and coupling strength also have a highly significant effect of electrode (p < 0.001). Prevalence has no significant effect of electrode (p = 0.30) and coupling strength has a weak effect of electrode (p = 0.04) that does not remain significant after false discovery rate correction. The frequency of the upper phase coupling mode is higher centrally compared to frontally but decreases more modestly centrally, with the frequency decreasing slightly from about 14.3 Hz at 10 years to about 14.1 Hz by 80 years. Frontally, the frequency decrease is more pronounced, from about 13.7 Hz at 10 years to ∼12.7 Hz by 80 years. The preferred phase of the upper coupling mode shifts leftward with age and is consistently closer to the trough frontally. Centrally, the preferred phase shifts from the peak of the SO (∼0.1 rad) at 10 years to the rising phase (∼−0.5 rad) by 80 years. Frontally, the preferred phase shifts from the falling phase (∼0.4 rad) at 10 years, reaches the peak by ∼60 years and continues toward the rising phase reaching −0.1 rad by 80 years. The coupling strength shows a very small but significant age-related increase through midlife that is larger frontally, followed by a slight decline. Centrally, the coupling strength increases from about 0.0133 at 10 years to about 0.0143 by 50 years. Frontally, the coupling strength increases from about 0.0126 at 10 years to the central coupling strength of about 0.0143 by 50 years. Centrally, the upper mode becomes less frequency dependent with age, decreasing from a mode tilt of ∼0.28 rad at 10 years to ∼0 rad by 80 years. Frontally, the frequency dependence follows a moderate quadratic age-related change, increasing from ∼0.2 rad at 10 years to 0.3 rad at 40 years, after which it begins to decrease. The prevalence increases with age at both electrodes, with no significant electrode difference, from about 68% at 10 years to about 86% by 40 years, after which it levels off near 88%.

### The lower phase-coupling mode weakens and drifts from the SO trough with age

The lifespan dynamics of the lower phase coupling modes are illustrated in the second row of Figure 6B and the model statistics are summarized in Table 2. The models for frequency, preferred phase, coupling strength, and frequency dependence, as well as for the proportion of subjects expressing the lower mode, all have a highly significant effect of age and electrode (p < 0.001), except for the preferred phase, which does not have a significant effect of electrode, prevalence, which has a significant effect of electrode (p<0.01), and frequency dependence, which has a weaker effect of age (p = 0.03) that does not remain significant after false discovery rate correction, and no significant effect of electrode. The frequency of the lower mode is higher centrally and shows different dynamics compared to the frontal electrode. Centrally, the frequency of the lower mode slightly increases from about 7 Hz at 10 years to ∼7.5 Hz at 40 years, after which it begins to decrease, reaching ∼6.4 Hz by 80 years, while frontally, the frequency stays relatively stable at about 6 Hz throughout 10 to 80 years. The preferred phase of the lower coupling mode shifts from the trough (∼±π rad) at 10 years toward the falling phase, reaching ∼2.2 rad by 80 years, with no significant effect of electrode. The coupling strength is higher frontally and shows an age-related decrease, with the decrease being more pronounced centrally. The coupling strength starts at about 0.014 at both electrodes and decreases to about 0.0125 frontally and 0.011 centrally by 80 years. The lower modes show little frequency dependence, with the tilt near 0 rad. The tilt appears to increase slightly with age to about 0.2 rad by 80 years. However, the effect of age is only nominally significant (p = 0.03) and there is no significant effect of electrode. The lower mode is most prevalent at younger ages and is consistently more prevalent centrally, with ∼90% of 10-year-olds having the mode, decreasing to about 65% by 80 years. Expression at the frontal electrode follows similar trends but is slightly lower by about 5%.

### Transient event heterogeneity may transcend traditional demographic categories

Previous work demonstrated marked heterogeneity of SOPH signatures within a small, triaged control group^40^, suggesting the potential for these spindle-like activity-based signatures to serve as individualized biomarkers of neural activity. We next examine the generalizability of these findings in this much larger population. Figure 7 explores heterogeneity in the context of increasing demographic restriction. Here, we show two groups taken from the age bins centered on 10 and 50 years (age: 10.5 ± 1.1 years, 50.1 ± 1.6 years) along with the mean SOPH for each group. Each group is demographically heterogeneous, composed of both male and female subjects across the different race/ethnicity categories. At the highest level, there are clear differences between age groups, with the older group expressing more modes with higher density overall. However, even after segregating by age, there are clear intersubject differences in mode features, and perhaps most distinctly, the number of modes expressed within each subject, even among subjects with the same race and sex. Further isolating subjects of the same age, sex, and reported race (pink boundary), stark structural differences in signatures remain. Finally, for each group, we also highlight (teal boundary) a pair of monozygotic twins (11 and 51 years) of the same age, race, and sex. In contrast to the large variations observed across the rest of the subjects in the cohorts, each set of identical twins shows striking similarities in both the classes expressed and the features within each class. Overall, these results suggest that individual differences in brain state may transcend even narrow traditional demographic categories. Moreover, the twin examples suggest the possibility of a genetic contribution to many properties of this high-dimensional signal. This finding is consistent with the demonstrated heritability of spindle features^6,55–57^ and EEG power spectra^9,58,59^, highlighting the importance of subsequent formal genetic studies of multi-class spindle-like activity.

**Figure 7:**
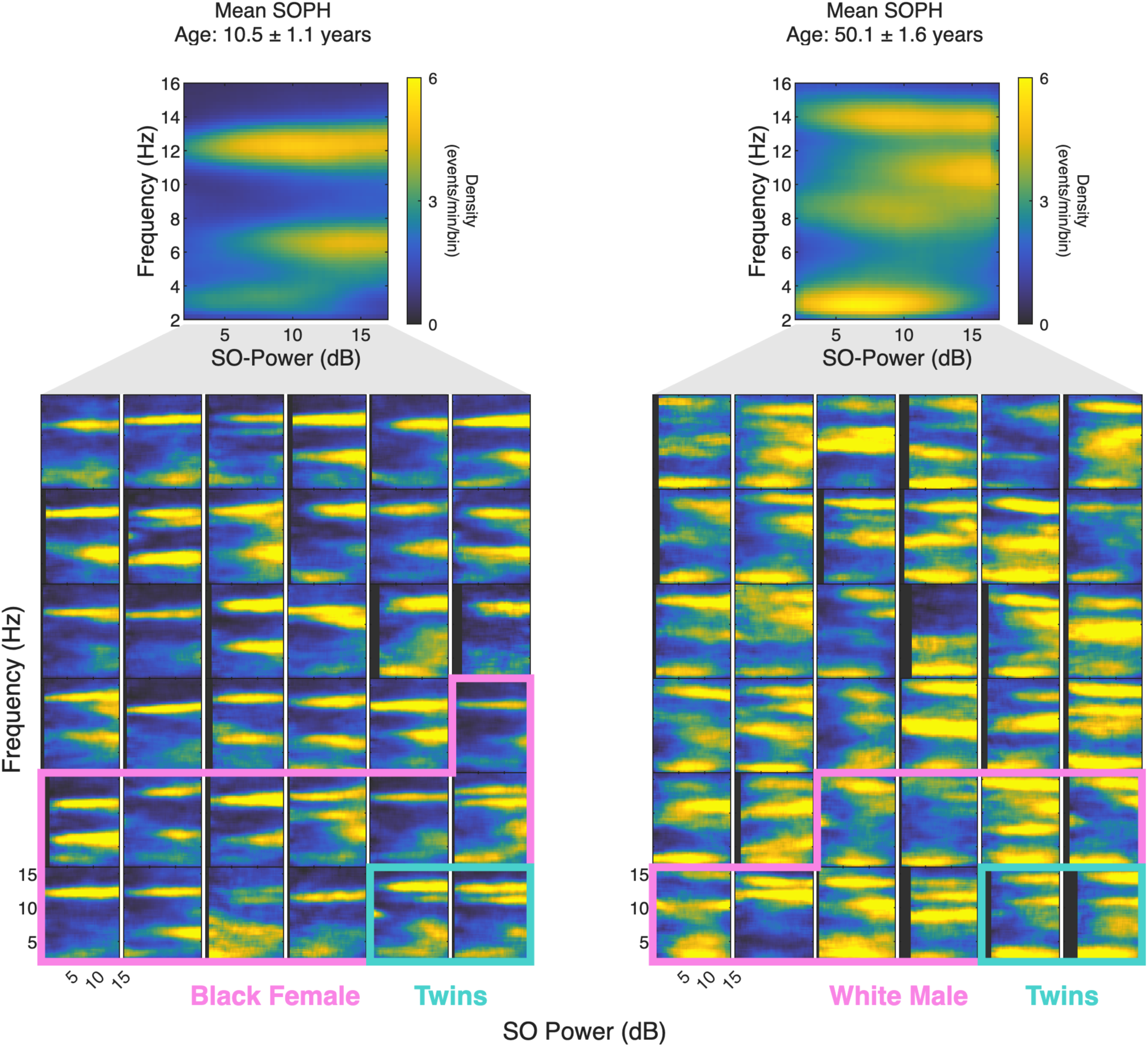
Individual heterogeneity in transient-oscillation class expression can transcend traditional demographics. Population mean SOPHs, while useful for identifying larger trends, can obscure the substantial heterogeneity in class expression across individual subjects. For two 5-year age bins centered at 10 and 50 years, we show the SOPHs of 36 randomly selected subjects from each bin, along with the mean age and group-average SOPH. The bins included 37 and 81 subjects, respectively. Across individuals, the density, frequency, and SO power of the classes vary considerably, and subjects also differ in which classes they express, neither of which is evident in the average. Even within a single demographic group (Black females on the left, White males on the right, outlined in pink) there is substantial heterogeneity in class expression. Within each of these groups we highlight a twin pair of the same race and sex, aged 11 and 51 years for the younger and older bins respectively, outlined in teal. The twins’ SOPHs appear more similar to one another than to those of the surrounding individuals, suggesting a potential genetic or shared-environmental basis for event class properties.

### Variation in event class expression provides a framework for phenotyping

Within the demographic cohort results (Figure 7), perhaps the most evident dimension of heterogeneity at the individual level is the variability in the expression of modes. We next explore the distribution of combinations of expressed classes and define individualized phenotypes that characterize subpopulations with similar neural states during sleep. For each subject, we identify the modes detected within the parameterized SOPH at a given electrode and then group subjects based on the combination of modes expressed. In Figure 8, we show the mode combinations that reach at least 15% prevalence in any age decile, for both the central (A) and frontal (B) electrodes. All remaining subjects are pooled into a single “All Others” group. A complete age decile distribution of all modes can be found in Supplemental Figure S1. For the subjects within each mode combination category, we show a scatter plot of mode frequency and sleep depth, as well as the group average SOPH. To quantify how phenotype prevalence changes across the lifespan, we fit a multinomial logistic regression that models the probability of each phenotype as a polynomial function of age. We show the median along with the 95% confidence intervals obtained from a family-level bootstrap.

**Figure 8:**
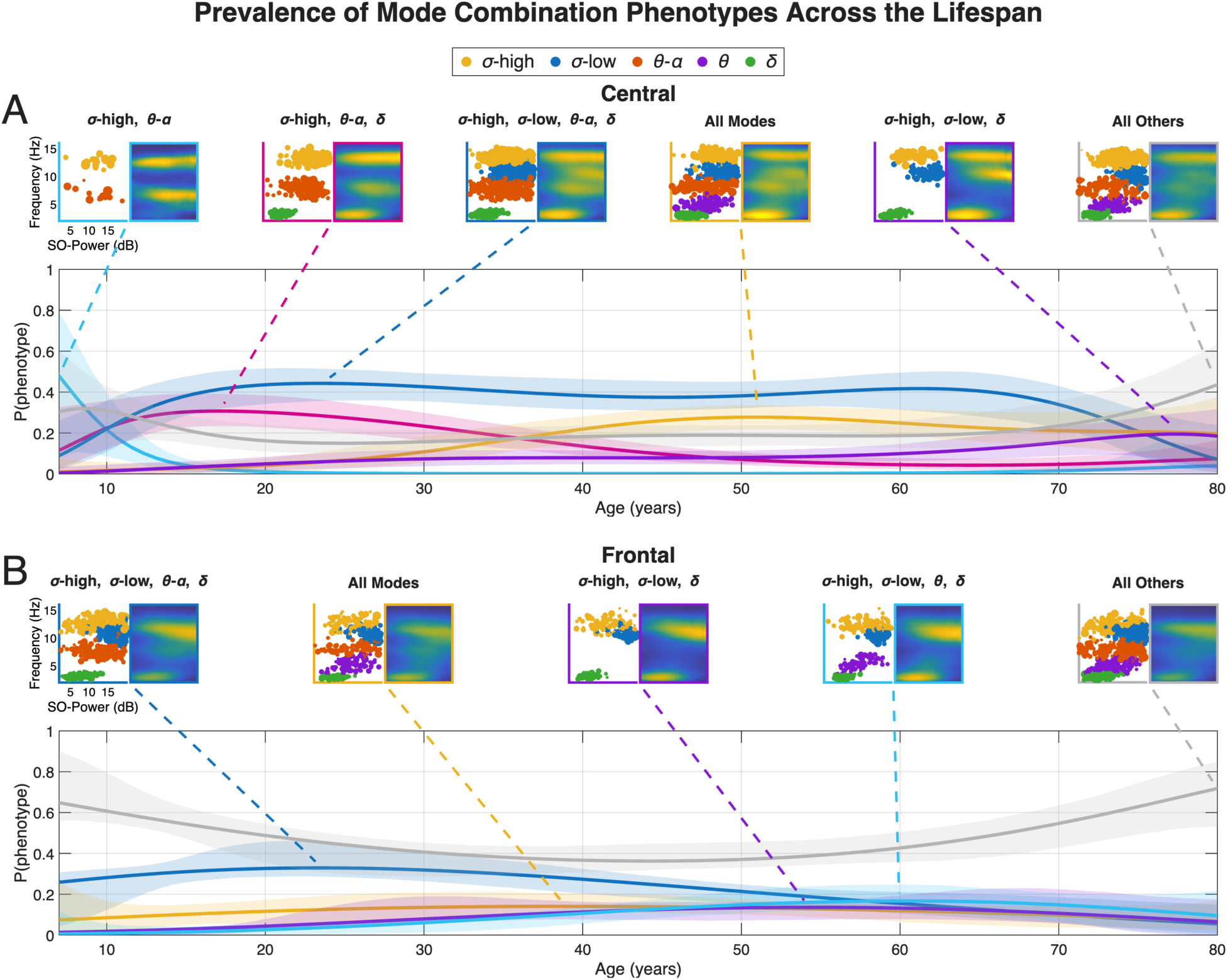
Event class expression as an individualized phenotype. From the previous analyses (Figure 7), it is clear that different subjects express different combinations of transient-oscillation classes. To explore the prevalence and dynamics of these potential individualized phenotypes, we categorized subjects by observed mode combinations in both the **A.** central (C3) and **B.** frontal (Fpz) electrodes. At the top of each panel, we show all mode combinations that reached at least 15% prevalence in at least one age decile. For each mode combination, we show the observed mode distribution in the frequency/sleep-depth space, with the individual modes colored by their assigned class (yellow: sigma-high, blue: sigma-low, orange: theta-alpha, purple: theta, green: delta), as well as the group average SOPH for all subjects at all ages who expressed that combination. All remaining subjects are pooled into a single “All Others” group, for which we likewise show the distribution and the average SOPH. To quantify how individualized phenotype prevalence changed across the lifespan, we fit a multinomial logistic regression that models the probability of each individualized phenotype as a polynomial function of age, and we plot these probabilities across age for every retained individualized phenotype and for the “All Others” group. The shaded bands are 95% confidence intervals obtained from a family-level bootstrap (whole families resampled), taken as the 2.5th and 97.5th percentiles of the bootstrap distribution. On each resample the polynomial order was re-selected, so the bands also reflect uncertainty in the chosen functional form in addition to sampling variability. For both electrodes, the probability over age of the top phenotypes was consistent with the overall age-related changes in the individual modes, with the frontal electrode showing greater heterogeneity indicated by the higher probability of the “All Others” group. Notably, no single top phenotype accounted for more than 0.5 probability at any age, highlighting heterogeneous class expression among individuals within the same age range.

Centrally (Figure 8A), the combination distribution generally reflects the patterns observed in the population analyses of lifespan trends (Figures 3 and 4). For the majority of the lifespan, from about 18–75 years, the most common combination is sigma-high, sigma-low, theta-alpha, and delta at about 0.45. This is consistent with the strongest modes observed in the population means (Figure 2). The two-mode combination of sigma-high and theta-alpha is maximal at the youngest ages (∼0.4 at 7 years), and the sigma-high/low, delta combination is maximal at the oldest ages (∼0.2 at 75 years), highlighting the progressive reduction in theta-alpha prevalence. Subjects with all modes appear at around 25 years, plateauing at approximately 0.3 by 50, reflecting the increasing prevalence of modes in theta with age. “All Others” combinations remain at roughly 0.2 throughout the lifespan, suggesting heterogeneity beyond these most common phenotypes. This reflects the previous observations that subjects of the same age can commonly differ in the combinations of classes they express. Moreover, it appears that late life is characterized by increased heterogeneity in the combinations of classes expressed. By 75 years, no single combination appears to strongly dominate, with many combinations contributing similar probabilities and an increase in the probability of the “All Others” group, which contains several other class combinations.

There is greater heterogeneity of expressed mode combinations frontally (see Figure S1). The “All Others” category is the most prevalent frontal phenotype at nearly every age (∼40–70%). While the most common central combination accounts for ∼40% of subjects in most age deciles, the most common frontal combination typically accounts for only 15–35%, with the remainder distributed across many combinations. This likely reflects in part the reduced separation between the sigma-high and sigma-low modes frontally, as well as a substantial subset of subjects for which a sigma-low mode (encompassing frontal slow spindles, see Discussion) is expressed without a detected sigma-high mode.

These results suggest that these different combination-based phenotypes can generally account for the mode structure observed cross-sectionally in the population, while also providing quantitative evidence that heterogeneity of class expression beyond aggregate trends is robustly present at the individual level. Future work exploring the environmental and genetic factors governing these differences will be critical for understanding the underlying biological and functional significance of these phenotypes.

## Discussion

This study establishes transient oscillatory event classes as data-driven, descriptive features of human sleep, which exhibit stereotyped dynamics throughout life, while retaining marked heterogeneity at the individual level. Overall, these results suggest that beyond changes in standard EEG band power, multi-class transient oscillation dynamics can serve as an important feature set with potential linkages to key mechanisms and networks involved in development, maturation, aging and related disorders, expanding on evidence suggesting broader classes of transient oscillatory activity^34,35,37,38,40^. Moreover, the ability to observe many features from multiple event classes with dynamics that differ across the lifespan provides a high-dimensional framework for identifying critical points and milestones in developmental and aging trends. By analyzing event class dynamics as a function of age, we can identify both distinct and gradual neural changes, offering valuable insights into the mechanisms and functionality of each class. We now highlight key dynamic milestones in terms of early (∼10–25 years), middle (∼25–50 years), and later (∼50–70 years) life.

### Early life (10-25 years) changes

#### Sigma-high dynamics in early life confirm existing studies of traditional fast spindles

The dynamics observed in the sigma-high mode directly reflect the results of numerous studies of traditional fast spindles, in particular, the clear rise in fast spindle frequency during childhood through adolescence^5–8^. Furthermore, the sigma-high frequency and density age-related dynamics directly parallel work from Purcell and colleagues^6^ using a sweep of wavelet detectors over a large frequency range across the lifespan. This is important because it suggests that many macro-scale findings from traditional spindle analyses using ad hoc frequency bounds and detection thresholds are also observed with our more general approach. In contrast, sigma-low is relatively stable during this period and has low prevalence centrally. Moreover, our results suggest that previously described slow spindles are a combination of classes that include that sigma-low (see later section for further discussion).

#### Frequency increase in theta-alpha provides a new potential developmental biomarker

We identified an increase in the frequency of the spindle-like activity classes in the theta-alpha and theta ranges which was most pronounced starting in early life. The theta-alpha event frequency increased from ∼7 to 7.8 Hz from the youngest subjects (minimum age: 7 years) to about 25 years of age. The frequency continued to increase to 8 Hz until about 40 years, but at a slower rate, suggesting that theta-alpha activity is particularly plastic during the transition from childhood to early adulthood. Additionally, the theta event frequency increased from ∼5 Hz at 10 years to ∼5.4 Hz centrally and ∼5.6 Hz frontally by 30 years. These findings provide a concrete electrophysiological mechanism underlying previous studies showing a shift from theta to alpha spectral power from 3–24 years^60^ as well as age-related decreases in theta power^5,7,61^. Theta-alpha and theta activity coupling near the slow-oscillation trough was also similar to reports of temporal theta–slow wave activity, a proposed biomarker of language and speech development observable as early as 24 weeks of gestation^62^. While it is not known whether these phenomena reflect the same underlying activity, these findings suggest that theta-alpha activity represents a potentially important marker of neurodevelopment from childhood into early adulthood.

In general, though the precise generating circuits and functional role of these theta classes remain unclear, theta activity has been strongly linked to memory processing across species^63,64^, and in humans, increased theta during memory encoding has been associated with sleep-related memory consolidation^65,66^ and memory reactivation^67^. Theta and alpha bursts have also been identified in REM^34,35^, and recent work has suggested a role for theta bursts in visual perceptual learning^68^. Cortical theta bursts have also been shown to coordinate with hippocampal sharp wave ripples and spindle ripples in humans^39,69^, and NREM theta bursts have been proposed to trigger the down-state–sharp wave ripple–slow oscillation/spindle cascade that supports long-term memory consolidation^38,69^, suggesting a critical role in memory consolidation that is different from that of classical spindles. Overall, these findings suggest that the age-related shift in theta-alpha frequency is a promising developmental feature for future study of memory-related changes during development.

#### Early-life frequency increase as a potential shared motif across oscillations that reflect brain maturation

Early life (10–25 years) was marked by an increase in oscillatory frequency across three spindle-like classes, with an increase of just under 0.5 Hz between ages 10 and 25 in both the sigma-high and theta classes, and a larger increase of about 0.8 Hz in the theta-alpha, suggesting a shared maturation-related motif. The increase in sigma-high frequency likely corresponds to previously reported developmental increases in fast spindle frequency^5–8^. To our knowledge, the developmental increase in theta-alpha spindle-like frequency observed here (discussed above) has not been previously reported during sleep.

Together, these findings indicate that early-life frequency acceleration is not limited to canonical spindles but may represent a broader feature of this developmental stage. The timing of the frequency increase coincides with a key period for neurodevelopmental changes, such as the increase in myelination that occurs during childhood and adolescence^70^. This is an important area for future clinical exploration given the disrupted myelination seen in neurodevelopmental disorders such as autism and schizophrenia^71^. The frequency of these oscillations may therefore potentially serve as an index of circuit development and maturation. For example, in fast spindles, frequency has been linked to cognitive development^72^ and has been proposed to reflect maturation and improved efficiency of corticothalamic networks, which are altered in neurodevelopmental disorders^20,72^. More broadly, given the role of spindles and theta activity in different aspects of memory and SO/sharp wave coordination^21,22,64^, the frequency increase might be a potentially important marker related to maturation of the circuitry underlying these processes.

#### The shift of sigma-high coupling to the peak of the SO may reflect refinement of timing important for memory consolidation during late childhood

Early life (10–25 years) was also marked by a rapid leftward shift in the preferred phase of the sigma-high coupling to the slow oscillation (SO), progressing from the falling phase of the SO (∼π/4 rad) around age 10, moving leftward toward the SO peak, reaching 0 rad by 35 years. Prior work has reported leftward shifts in fast spindle–SO coupling away from the SO peak during aging, which have been linked to age-related declines in memory consolidation^16,73^. There have also been reports of developmental changes in spindle and SO phase coupling between childhood and adolescence^7,54^, and increases in coupling strength have been associated with improvements in various memory tasks^54,74^. The present findings extend this literature by demonstrating that the leftward shift in coupling occurs throughout life, with a pronounced early-life leftward shift toward the peak of the SO. Interestingly, a leftward shift in coupling was observed across all classes, suggesting this may represent a general motif. However, the other classes never reached the peak of the slow oscillation.

Since precise coupling of fast spindles to the peak of the slow oscillation is thought to be critical for memory consolidation and synaptic plasticity mediated by increased calcium influx during cortical up-states at the SO peak^16,50–52^, the age-related shift in the sigma-high coupling observed here may reflect a developmental phase of spindle–SO coordination. In contrast to aging, where coupling shifts away from the peak are associated with impaired memory consolidation, the early-life shift toward peak coupling may signify refinement of spindle timing that supports increasingly efficient memory processing. Given that this developmental period is marked by substantial improvements in declarative memory^75^, future studies incorporating memory tasks will be important to determine whether the age-related shifts in spindle–SO coupling observed here are associated with declarative memory performance.

### Midlife (25-50 years) changes

#### Midlife was marked by the peak expression of two classes of activity

The transition from early to midlife (∼25–50 years) was marked by the strongest prevalence of two classes of activity, sigma-low and delta. The sigma-low class showed a spatial change in expression during this transition, with increased expression centrally. Sigma-low activity was prevalent frontally in about 90% of individuals from 20 to 80 years, but it did not reach peak prevalence centrally until midlife, with the highest proportion of individuals expressing this class around 40 years, after which central sigma-low prevalence plateaued. Because these events likely represent a subset of what has previously been grouped under slow spindles (discussed in a later section), their specific developmental dynamics have not been separately characterized in prior work. Given that the sequence and timing of nested oscillations are thought to provide windows for different forms of neural communication and plasticity^50,76^, and that sigma-low activity occurs closer to the slow oscillation trough, future work should determine whether these events serve functions distinct from the canonical memory consolidation processes typically attributed to fast spindles, such as later phases of sleep-related memory processing.

Delta activity steadily increased from childhood to midlife, where it was most prevalent, with the highest proportion of individuals expressing this class around 40 years, after which the prevalence plateaued. In contrast, broad delta power during sleep has been shown to decline beginning around adolescence^5,7,61^. Future studies will be important for understanding the relationship between the decrease in broad delta power and our observed increase in transient delta events with age. Prior work on delta activity provides several hypotheses regarding its potential relevance. The emergence of delta bursts across the lifespan contrasts with the decline in broadband delta power, which has been linked to cortical thinning trajectories and hypothesized to reflect synaptic downscaling processes^61,70,77,78^. One possibility is that reduced background delta activity makes discrete bursts more prominent. In addition, alterations in delta activity have been associated with changes in sleep pressure^79–81^ and reductions in time spent in deep sleep^17^, raising the possibility that delta bursts might reflect increasing sleep fragmentation across aging. Finally, recent work in rodents has identified a 2–5 Hz oscillation in the prefrontal cortex that couples prefrontal and hippocampal activity in anesthetized animals^37^, as well as a 2–4 Hz hippocampal oscillation coherent with nasal respiration^36^, suggesting that delta-range bursts may reflect additional processes whose functional role remains to be determined.

#### Midlife is characterized by a spatial change in the expression of sigma-high

We observed clear spatial changes in the proportion of subjects expressing sigma-high activity, characterized by a reduction in the proportion of individuals expressing sigma-high spindles at the frontal electrode. Specifically, while the proportion of subjects expressing sigma-high activity at the central electrode remained relatively stable from approximately 10 to 80 years of age, frontal expression declined, with the most pronounced decrease beginning around 45 years and continuing to late life. This age-related change in spatial expression is consistent with prior reports indicating that fast spindle activity becomes more posterior with age^8,82^. The theta-alpha events showed a similar, although less pronounced, age-related change in spatial expression, suggesting that this pattern may extend beyond fast spindle activity. Future work using high-density EEG and source localization will be informative for better understanding the extent of this spatial shift and potential functional segregation. One possible interpretation of this finding, particularly for the sigma-high, is that the change in spatial expression reflects functional refinement of corticothalamic circuitry that underlies spindles across adulthood, corresponding to age-related increases in the functional segregation of the precuneus, a region identified as a major cortical source of fast spindles^82–84^. In this context, the presence or absence of frontal sigma-high activity may serve as an index of functional specialization within spindle-generating networks.

#### Late life (50-70 years) is marked by a general uncoupling of transient events from the SO

In the transition from mid- to late life (∼50 years), we observed a continuation of the pronounced reduction in the coupling strength as well as the proportion of individuals expressing strongly coupled lower phase modes with age, and the continuation of the leftward shift in the preferred phase of the upper coupling mode, which brought it from the SO peak toward the rising phase. Prior work has reported aging-related leftward shifts in fast spindle–SO coupling that were linked to declines in memory consolidation and cortical atrophy in medial prefrontal regions^16,73^. Given evidence that precise spindle–SO coupling is important for effective memory consolidation^16,50–52^, the age-related uncoupling observed in late life may reflect disruptions in memory processing that accompany aging^16,85,86^. Although the specific functional role of the lower phase coupling mode remains unclear, it likely overlaps with the events observed in the theta-alpha and theta range. The slower decrease of coupling strength frontally may suggest a distinct or more resilient functional role for frontal oscillatory interactions in older age.

#### Frequency-dependent phase coupling reveals a novel axis of fast spindle changes across the lifespan

The upper phase-coupling mode, spanning approximately 10–16 Hz, exhibited strong frequency-dependent coupling in childhood that weakened with age at the central electrode. Specifically, the tilt of the upper phase-coupling mode decreased toward zero across the lifespan centrally, indicating a loss of frequency dependence, while remaining relatively frequency dependent from 20– 60 years frontally. In childhood, higher-frequency events preferentially coupled to the rising phase of the slow oscillation, whereas lower-frequency events showed progressively rightward shifts in preferred phase toward the falling phase. By adulthood, however, events in the upper phase coupling mode across the entire frequency range had the same preferred phase, independent of frequency. This suggests that there are differences in the coupling of spindles to the slow oscillation as a function of frequency.

Previous work has reported differences in fast spindle–slow oscillation coupling as a function of other factors such as sleep stage and spatial location^87–91^. Of particular interest are the previously reported two fast spindle subtypes, “early-fast” and “late-fast,” that preferentially couple to the rising phase and peak of the slow oscillation and are more prevalent in younger and older individuals, respectively^90,91^. Our findings suggest that these previously described subtypes may reflect a continuum within a single phase-coupling mode, captured by age-related changes in frequency-dependent coupling rather than representing distinct spindle classes.

This change in frequency dependence points to a previously unstudied axis of developmental change in which phase coupling becomes progressively less frequency dependent with age, potentially reflecting refinement or streamlining of coupling mechanisms across development. The presence of frequency-dependent coupling in childhood may indicate greater flexibility in how spindle activity engages different slow-oscillation phases, while the convergence toward a single preferred phase in adulthood may reflect increased temporal precision. The differing dynamics of the frequency dependence at the frontal electrode suggest potential regional differences in how coupling mechanisms mature, which may support distinct memory systems or stages of consolidation. High-density EEG studies will be important for determining whether these developmental changes in frequency dependence occur along a spatial gradient.

Finally, prior reports of increased spindle–slow oscillation coupling strength from childhood to adolescence^54,74^ (which we also observe) may be partially explained by the age-related reduction in frequency-dependent phase coupling observed here. Because coupling strength is sensitive to variance, unmodeled frequency dependence may artificially inflate variability and reduce apparent coupling strength. The persistence of frequency dependence at frontal sites suggests that additional sources of variance reduction may contribute to frontal coupling changes across development.

#### Class expression reveals brain activity-based individualized phenotypes

Grouping participants based on the combination of expressed classes reveals a set of distinct phenotypes, whose prevalence changes across the lifespan. Importantly, at every age, multiple phenotypes are observed, indicating that individual variability is not simply a byproduct of age-related trends. The factors underlying these phenotypes remain to be determined, but they could potentially reflect a combination of genetic, environmental, and age-related influences, requiring careful interpretation. In particular, it is important to understand the various factors contributing to the absence of a given class. For some subjects, a class may be present but fall below detection criteria or may have activity that is too diffuse or skewed to be captured by Gaussian-like basis functions. Alternatively, the activity may not be absent but rather expressed in a different spatial location. In cases where a class is truly absent, this may point to more fundamental differences in neural circuitry, raising the possibility that different phenotypes correspond to distinct genetic, physiological, or clinical profiles.

Given the amount of heterogeneity observed, even within traditional demographics, it is important to not jump to the conclusion that absence of a detected class implies any pathological state, similar to reports that ∼10% of the population does not express eyes-closed alpha^92^. Thus, further exploration of the implications of class expression phenotypes is warranted. Regardless, awareness of expression-based phenotypes is essential for future studies of spindle activity. Lack of control for these differences, especially in small cohort studies, could lead to spurious differences, simply due to the heterogeneity found even within demographically and age-controlled cohorts (Figure 7).

#### Classically reported “slow” spindles are likely a mix of multiple classes of activity

Our results suggest that traditionally defined ∼9–12 Hz “slow spindles”^6–8^ in the literature likely reflect a combination of multiple distinct spindle-like activity classes rather than a single homogeneous event type. Specifically, we identify three classes that plausibly contribute to prior reports of slow spindles: frontal sigma-high activity, which occurs throughout NREM sleep and across the lifespan; sigma-low activity, which occurs deeper in NREM sleep and shows an age-related spatial change to increased central expression during midlife (∼25–50 years); and theta-alpha activity, which becomes earlier in NREM sleep with age and is most prevalent at younger ages. All of these classes cross into the 8–12 Hz range at some point in the lifespan and are present at the frontal electrode. Yet, our analyses show these classes are distinct in their properties and lifespan dynamics.

This interpretation addresses substantial variability in how slow spindles have been defined and characterized across prior studies. Reported frequency boundaries for slow spindles vary widely, with Cox et al.^89^ noting that the demarcation between fast and slow spindles ranges from 12–14 Hz, and that the lower bound for slow spindles spans 8–12 Hz, overlapping with the alpha/theta range. Similarly, reports of phase coupling relative to the SO have been inconsistent. Some studies have described slow spindles as coupling to the falling phase of the SO^50,87,93,94^, whereas others have reported that both fast and slow spindles follow the down-state and couple near the SO peak, with only theta activity preceding the down-state^38,69,95^. In addition, the functional role of slow spindles remains less clearly defined than that of fast spindles, with mixed findings across studies^89,96–99^. In light of these differences, some authors have questioned whether slow spindles constitute a distinct class at all^22,95^. Distinguishing among event classes may help future work clarify the functional roles previously attributed to slow spindles, which may in fact reflect contributions from multiple oscillatory processes with potentially different roles.

#### Potential origins and mechanisms of transient events

The distinct frequency, sleep depth, and phase-coupling dynamics observed across transient oscillation classes suggest that each event class may serve different roles during sleep across the lifespan. Although the present study does not establish functional significance, this diversity of oscillatory dynamics raises the possibility that different classes contribute to distinct aspects of sleep-dependent processing. There is evidence from the literature that can provide insight into the potential mechanisms observed in this study. Classical spindles in the sigma range are known to arise from recurrent interactions within thalamocortical networks^10,22^. Prior work using human intracranial recordings^38^ has suggested that theta burst activity exhibits distinct corticothalamic dynamics from spindles, with theta bursts occurring more equally across cortex and thalamus and showing less temporal overlap between regions compared to spindles, which are more strongly expressed in the thalamus.

#### Limitations and extensions

In this study, we focused on transient oscillatory spindle-like activity recorded from a single central and frontal EEG channel in a large, cross-sectional dataset. Further studies on other data can augment our understanding in various ways.

While evidence suggests that mechanistically distinct generators may contribute to events within each class, the events themselves are defined on a purely morphological basis. A key question, then, is whether they reflect genuine activity from specific networks or merely robust detectable patterns arising from secondary effects of other EEG features. Thus, as is the case for all pure EEG analyses, there are some questions of interpretation. For example, given a fixed duration, a transient oscillation becomes increasingly ill-defined as the frequency decreases, because the number of oscillation periods becomes progressively smaller. In the EEG, a decreasing frequency also means an increase in the aperiodic noise, further reducing the signal-to-noise ratio. Furthermore, it is mechanistically important to make the distinction between an event such as a spindle with distinct initiation, maintenance, and termination phases, and a persistent oscillation that manifests as bursts due to fluctuations in amplitude. Future quantitative studies can attempt to resolve these differences by using statistical time-series models of EEG data with latent variables that allow for state-switching^100^. Through model selection, it will be possible to see whether the data are better described by discrete state switches or by stochastic variation in the oscillation amplitude. Conclusive determination of functional and mechanistic significance therefore requires careful consideration and interpretation. Regardless of the mechanisms, our current approach allows us to identify events through their morphological traits and show significant age-based dynamics in their properties.

While these analyses do not include neonatal or early childhood data (minimum age: 7 years), future work tracking the evolution of these event classes starting at birth through the first few years of life will provide key insights into neurodevelopment and early childhood critical periods. Additionally, the development of large-scale longitudinal datasets with multiple data points per subject throughout the lifespan will be essential for direct measurements of individual trajectories and assessing the relationship between variability in individual trajectories and clinical outcomes. As these are extremely costly and difficult experiments to undertake, support for long-term longitudinal sleep studies and large-scale retrospective clinical database analysis will be important for advancing the field.

Our analyses focused on characterizing waveform classes in low-density scalp EEG, from which neither the underlying topological distribution of events nor the circuit mechanisms responsible for their generation can be directly inferred. Future work using high-density EEG can better characterize event classes across a spatial gradient^8,88,89,101^ and source localization could provide more detailed putative cortical distributions. Ultimately, extensions to human intracranial recordings as well as animal model studies with local field potential and spiking data will be critical for elucidating the roles and sources of these different classes.

## Methods

### Dataset description

The data analyzed in this work from the Cleveland Family Study^42,43^ were obtained from the National Sleep Research Resource^42^ (sleepdata.org). The Cleveland Family Study is a cross-sectional study spanning ages from 7 to 89 years, with a mean age of 41.4 years, 55% female, 56% Black, 41% White, and 3% Other. EEG data were sampled at 128 Hz, and the C3 and Fpz electrodes were re-referenced to linked mastoid. Of 735 total subjects, 5 were excluded because a SOPH could not be computed from their recording (insufficient data), and an additional 26 were excluded after visual inspection due to corrupted SOPHs from excessive line noise, leaving 704 subjects for analysis. For this subset of 704 subjects, the demographics are similar to the full dataset, with a mean age of 41.5 years, 54.5% female, 55.5% Black, 41.5% White, 3.0% Other.

### Artifact detection

Artifact rejection was performed in the time domain using an iterative outlier removal procedure^40,102^. Briefly, the raw data were filtered into higher-frequency (35 Hz to Nyquist) and lower frequency (0.1 Hz to Nyquist) components. In each band, the amplitude envelope was smoothed (2 s), log-transformed and detrended with a 300 s moving median. Any data with a robust z-score (median and median absolute deviation) above 5.5 was marked as artifact and removed. The z-score was recomputed and the process iterated until no data exceeded the threshold. Ten-second windows whose spectral slope over 1–55 Hz exceeded −0.5 were also marked as artifact.

### Detection of spindle-like transient oscillatory events

For each subject, spindle-like transient oscillatory events in the 2–16 Hz frequency range were automatically detected using the DYNAM-O toolbox^40^ with default parameter settings. Briefly, the approach relies on the principle that transient oscillatory activity in the time domain, which is characteristic of spindle activity, appears as time-frequency peaks in the time-frequency domain. To account for the aperiodic background, each spectrogram was divided at every frequency by a static baseline spectrum, the 2nd percentile of power across the scored period of the record excluding artifacts. DYNAM-O operates in the time-frequency domain, using a modified watershed algorithm to detect time-frequency peaks (which correspond to transient oscillatory activity) across a broad frequency range from the full night spectrogram of activity, and extracts the peak properties (such as the central time, frequency, amplitude, and duration) along with estimates of sleep depth and slow oscillation phase. DYNAM-O conservatively retains only the peaks resolvable under the spectral estimators, thereby reducing any bias caused by ad hoc noise rejection thresholds. Thus, DYNAM-O aims to identify a more objective and holistic set of spindle-like transient oscillatory activity throughout the night, compared to traditional threshold-based methods.

We summarize the activity of tens of thousands of these events throughout the night by creating two different mappings of the data at different temporal scales. The *slow oscillation power histogram* (SOPH) is defined as the mapping of TF-peak density as a function of frequency and sleep depth, where sleep depth is measured through slow oscillation (0.3–1.5 Hz)^103^ power. The *slow oscillation phase histogram* (SOΦH) is defined as the mapping of the probability of observing a TF-peak given frequency and the phase of the slow oscillation. Together, the power and phase histograms provide an individualized signature of TF-peak activity across multiple classes of events and multiple time scales.

### Ǫuantitative modeling of event class properties

#### Mode parameterization and classification

TF-peaks with similar properties will naturally cluster within a parameter space, forming modes representing different classes of spindle-like events. These modes are clearly observable in the SOPH and SOΦH as regions of enhanced density/event probability. The combination of the features of these modes represents each participant’s individualized signature. We used the DYNAM-O toolbox to apply a parametric modeling approach to characterize the properties of the event classes observed within each subject. Herein we describe the overall process in brief; see He et al.^104^ for a detailed overview of the underlying algorithm.

For both power and phase histograms, the model decomposes the histogram, *H*, into a set of *N* mode basis functions,*m*_1…_*m_n_* and an underlying baseline function, *B*:

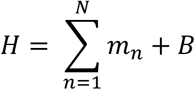

When fit to a given SOPH/SOΦH, the parameters of the fit modes serve to quantify the features of each class of events observed within the data. The SOPH was modeled as a sum of a set of rotatable two-dimensional Gaussian basis functions and a planar baseline. Similarly, the SOΦH was modeled as the sum of a set of von Mises (Gaussian in the circular domain) functions over a fixed cosine-tuned baseline. Each mode contains several parameters including amplitude, location in the histogram, standard deviations in both dimensions, and axial rotation.

For the SOPH, the location parameters define the mode center in the frequency-sleep depth (SO power) space, the amplitude defines the maximum mode density, the standard deviations define the extent/bandwidth of the mode, and the rotation defines the rate of frequency change with sleep depth. For the SOΦH, the location parameters define the mode center in the frequency-preferred phase (SO phase) space, the amplitude defines the maximum mode coupling strength, the standard deviations define the extent/bandwidth of the mode, and the rotation defines the frequency dependence of the preferred phase. Given a central frequency for each observed SOPH mode as defined above, we estimate that mode’s preferred phase as the phase of maximal coupling in the corresponding SOΦH model fit at that frequency. It is important to note that this value is distinct from the preferred phase of the phase coupling mode, which describes the macrostructure of event coupling across frequencies.

The power mode classes were identified through a data-driven clustering approach, described in the next section. The phase coupling classes were defined based on frequency. We used the ranges of 10–16 Hz and 4–10 Hz to define the upper and lower modes, respectively. Phase modes outside both ranges were not assigned to a class and were excluded.

Before classification, modes were required to have a center frequency between 2 and 16 Hz. For power modes, the SO power was also required to exceed 2 dB. We additionally censored modes whose tilt (power modes) or frequency dependence (phase modes) was large in absolute value, retaining only power modes below 0.15 rad and phase modes below 1 rad. This kept the included modes approximately axial with respect to the primary axis of their distribution.

### Data-driven clustering of transient oscillation modes

Given the set of parameterized modes across a population and multiple electrodes, it remains essential to separate the data into distinct classes of modes. Given that a major motivating factor in this work is to minimize a priori assumptions, rather than define frequency bands, we used an unsupervised clustering approach to provide data-driven classification of transient oscillation modes based on mode frequency and sleep depth. To do so, we adapted a standard Gaussian mixture model (GMM) approach to account for nonlinear frequency dynamics with age, data noise, and individualized signatures that could be masked by large population variability. Class number and properties were also chosen in an unsupervised manner. Clustering, including the age remapping described below, was performed separately for each electrode.

We first removed nonlinearities in mode frequency as a function of age through a remapping procedure. For the remapping, we computed the smoothed number of modes per subject as a function of age and mode frequency, in sliding 10-year age windows (1-year step, each window was adjusted, if necessary, to contain at least 80 subjects to ensure sufficient data for estimates throughout). In this density, distinct classes of modes will appear as continuous ridges evolving across age. All observed ridges were identified through peak tracking (MATLAB findpeaks()), and the identified continuous ridgelines were used to create a monotone piecewise-cubic mapping, which mapped the ridge frequencies at each age onto their frequencies at the median age, so that all ridges had a constant frequency across age. Only frequency was remapped, and this mapping was then applied to all modes from all subjects. We then fit a full-covariance Gaussian mixture model with an added uniform background component that absorbs noise and artifact modes^105^. Given that the separation between a subject’s modes is often far less than the population variability, it was important to look at this as a problem of clustering linked sets of modes from individuals rather than finding clusters in the pooled population space. To address this, we constrained the GMM by matching. That is, each subject expresses each class at most once, with each class possessing an independent Bernoulli probability rather than a mixture weight. Remaining noise modes are defined as a Poisson background, uniform over the observed feature range, with the rate estimated alongside the class parameters. A subject’s modes therefore compete for classes, as in a Gaussian mixture under cannot-link constraints^106^, and the subject likelihood sums over all assignments in which each class takes at most one of the subject’s modes and the remaining modes are background^107^. The model was fit by expectation-maximization^108^. Each fit was initialized from a k-means partition of the z-scored features^109^.

We assessed goodness-of-fit using the integrated completed likelihood criterion (ICL)^110^. In order to be robust to local minima, the result was chosen as the minimum ICL of five restarts of the fit. The number of classes, K, was selected by sweeping K = 3–7 and choosing the elbow of the ICL curve using the Kneedle algorithm^111^. K = 5 was selected at both electrodes. Each subject’s modes were then assigned to their most probable configuration; modes assigned to the background were left unclassified.

### Lifespan dynamics regression analysis

We modeled each parametric feature of each oscillatory class as a function of age. Modes from the two electrodes (central, C3; frontal, Fpz) were analyzed jointly with electrode included as a within-subject factor, and classes were modeled separately. For each class and feature, the feature was regressed on a polynomial in age and adjusted for sex, electrode, race (white vs. non-white), body-mass index (BMI), and apnea–hypopnea index (AHI), with age × sex and age × electrode interactions and a subject random intercept accommodating the multiple modes and both electrodes contributed by each subject. An identical covariate structure was used for all features. Because the Cleveland Family Study enrolled families (704 subjects in 144 families), the Gaussian and Gamma models also included a family random intercept. The von Mises model supports only one random intercept, which was used for subject. Its standard errors were instead computed treating each family as one independent unit.

The response distribution was chosen to match each feature. Frequency, slow-oscillation power, density, and frequency dependence were fit as Gaussian linear mixed-effects models (fitlme). Coupling strength was fit as Gamma generalized linear mixed-effects models with a log link (fitglme). Preferred slow-oscillation phase (of the phase modes, and of the power modes at their center frequency), a circular variable, was fit with a von Mises mixed-effects regression in which each observation is von Mises about the linear predictor and each subject carries a von Mises–distributed random angular offset. The offsets were integrated out analytically and the resulting marginal likelihood maximized by expectation-maximization, with the regression coefficients updated by iteratively reweighted least squares and a weakly informative half-Cauchy prior (scale 8) on the between-subject phase concentration to prevent boundary degeneracy. Mode tilt, although angular, is confined to a narrow range far from the ±π discontinuity, where a von Mises model is weakly identified and less powerful than a linear one. Mode tilt was therefore modeled as Gaussian.

The polynomial order in age (from 0–2) was selected by the Akaike information criterion (AIC)^112^ for all models (Gaussian, Gamma, and circular), which were fit by maximum likelihood (Gaussian), Laplace-approximated maximum likelihood (Gamma), or penalized maximum likelihood (von Mises). Age entered as raw powers (in years) in the Gaussian and Gamma models and as an orthogonal polynomial basis in the von Mises models. The age effect for each feature was evaluated with an omnibus Wald test of all age-related terms, including the age × sex and age × electrode interactions; sex and electrode were tested analogously over their main effect and age interactions, and race, BMI, and AHI as single coefficients. Tests were F-referenced, except the Gamma covariate tests (χ²), and the von Mises tests used a covariance clustered by family. Goodness of fit was summarized by marginal R² for the Gaussian and Gamma models^113^ and by a circular R² (one minus the ratio of the summed 1 − cos residual angle to the summed 1 − cos deviation from the circular mean) for the von Mises models. Regression estimates were obtained by prediction at the population (zero random-effect) level for each electrode, with sex and race held at their most common category, AHI and BMI held at their means, and 95% confidence bands derived from the coefficient covariance. P-values are reported without correction for multiple comparisons, because each test addresses a single feature’s age trajectory. Sex, race, AHI and BMI are adjustment covariates. At a 5% false discovery rate (Benjamini–Hochberg) across all 319 tests, 62 of the 64 age and electrode effects with p < 0.05 remain significant. The two that do not are upper-mode coupling strength by electrode (p = 0.04) and lower-mode frequency dependence by age (p = 0.03).

Class prevalence was modeled at the subject level, over all subjects with an SOPH at that electrode, including those with no modes of the class (704 subjects). For each class, a binary indicator of expressing at least one mode of that class at a given electrode was regressed on an age polynomial with additive effects of sex, electrode, race, BMI, and AHI by binomial logistic regression (fitglm). Subject and family random intercepts were omitted because near-complete separation over portions of the age range drives a binomial mixed model to saturation. Inference accounting for the dependence between a subject’s two electrodes and among relatives was instead obtained from a family-level bootstrap (5,000 resamples of whole families) stratified by decile of family mean age, providing percentile confidence bands and Wald tests computed from the bootstrap covariance. The polynomial order in age was selected by AIC on the full sample, as above, and held fixed across bootstrap replicates.

### Modeling class expression-based phenotype probability over the lifespan

To define individualized phenotypes, each participant with at least one classified mode on a channel (702 subjects at each electrode) was characterized by the combination of classes they expressed, as determined from the modes identified by the parametric models. Combinations were defined as binary strings over the five power-mode classes, with a 1 reported if one or more modes of that class were identified and a 0 if none were found. Within each channel (C3, Fpz), a combination was retained as its own outcome category if it reached at least 15% prevalence in at least one age decile, where deciles were equal-count bins of the participant age distribution. All remaining combinations were pooled into a single “All Others” category. To describe the change in phenotype composition across age, we modeled the prevalence of these phenotype categories as a function of age using nominal multinomial logistic regression (MATLAB mnrfit), with phenotype category as the outcome and a polynomial in age as the predictor, fit separately for each channel. Polynomial order was selected by AIC, up to a maximum of fourth order. Predicted phenotype probabilities were then estimated across age. We obtained 95% confidence intervals from a family-level bootstrap with 5,000 resamples of whole families, taken as the 2.5th and 97.5th percentiles of the bootstrap distribution. Within each replicate the polynomial order was re-selected, so that the bands reflect uncertainty in the chosen functional form in addition to sampling variability.

## Acknowledgments

REB is a Research Health Scientist at VA Boston Healthcare System. The contents of this work do not represent the views of the U.S. Department of Veterans Affairs or the United States Government. The Cleveland Family Study (CFS) was supported by grants from the National Institutes of Health (HL46380, M01 RR00080-39, T32-HL07567, R01-46380). The National Sleep Research Resource was supported by the National Heart, Lung, and Blood Institute (R24 HL114473, 75N92019R002).

