## Supplemental Figures for "Multiple classes of spindle-like oscillations reveal stereotyped lifespan dynamics amid individual variability"

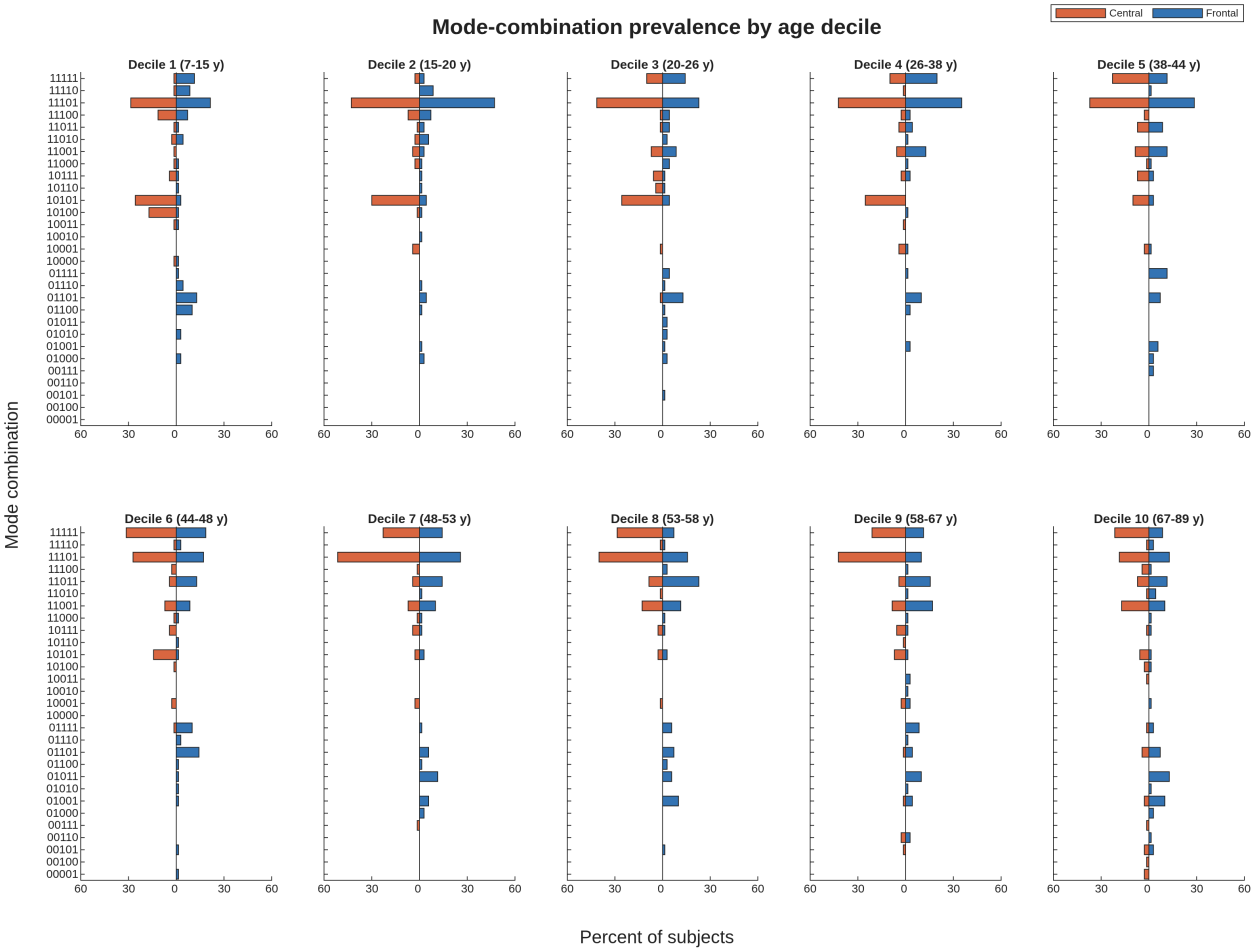


**Figure S1: Full breakdown of class expression-based phenotypes.** Since we observed differences in the expression of the different spindle-like classes across individuals within the same age range, we grouped individuals into phenotypes based on which combinations of modes/ individualized classes they expressed. In Figure 8, we showed the age-related trajectories of the phenotypes that had at least 15% prevalence in at least one decile and grouped the remaining phenotypes into the "All Others" category for simplicity. Here, we show the full breakdown, with the prevalences (percent of subjects) of each phenotype for each decile for the central (orange) and frontal (blue) electrode. Because there are 30 phenotypes across the deciles, we refer to the classes using binary notation. The binary notation has five digits, with each digit corresponding to one of the five spindle-like classes in descending order of frequency. A 1 or 0 indicates whether that phenotype expresses the corresponding class. For each decile, the top ranked phenotypes generally followed the age-related changes in prevalences of the individual classes, with the frontal electrode occupying a larger number of phenotypes compared to the central electrode.
